# Autism Mutations Preferentially Target Distributed Brain Circuits and Cell Types Associated with Sensory Integration and Decision Making

**DOI:** 10.1101/2025.08.13.670113

**Authors:** Jiayao Wang, Jonathan Chang, Andrew H. Chiang, Dennis Vitkup

## Abstract

Autism spectrum disorder (ASD) is a highly heritable psychiatric condition characterized by difficulties in social communication and stereotypic, repetitive behaviors. Genetics studies have discovered many dozens of genes with causal roles in autism, and functional analyses have demonstrated that ASD-associated mutations affect a diverse range of brain regions and cell types. However, the precise mechanisms by which these genetic alterations lead to autism-related phenotypes remain unclear. Psychiatric cognitive and behavioral traits are believed to arise from dysfunction of specific brain circuits formed by anatomically and functionally connected brain areas. To identify the circuits and cell types primarily affected by ASD mutations, we developed an unbiased approach, GENCIC, which computationally integrates genome-wide genetic data with a brain-wide spatial mouse transcriptome and an anatomical mesoscale connectome. Applying this approach to ASD reveals a convergence of mutations on cohesive brain circuits that are enriched in long-distance connections and span both cortical and subcortical brain structures. Furthermore, our analysis of brain-wide single-cell spatial transcriptomics shows that the heterogeneity of circuit structures affected in ASD is matched by the substantial diversity of strongly impacted circuit cell types. Notably, the implicated circuits and cell types play a central role in the integration of multimodal sensory and emotional information and in decision-making based on this information. We also find that different circuit structures exhibit distinct vulnerability patterns that correlate with cognitive phenotypes in ASD. Overall, our study demonstrates how ASD-related genetic mutations impact multiple levels of brain organization, ultimately disrupting functional circuits that drive core autism-related behaviors.

## Introduction

Autism is a common and highly heritable neurodevelopmental disorder^1^. Autism spectrum disorder (ASD) is typically associated with difficulties in communications, social interactions, and restricted and repetitive behaviors^2^. ASD also frequently leads to problems in motor skills and cognitive abnormalities. In the last decade, large-scale genetic studies uncovered many dozens of genes with strong causal links to autism^3–7^. Based on the growing sets of implicated genes, various functional and network-based approaches revealed specific molecular and cellular processes that are perturbed in ASD^8^. The strongly affected processes include synaptic communications, cellular signaling and neural development, and chromatin modification and regulation^9–11^. Analyses of expression data across brain regions and developmental periods found that ASD mutations impact multiple cortical and subcortical brain areas and diverse neural cell types^8,12,13^. Despite rapid progress in ASD genetics, it is currently not well understood how genetic perturbations at the molecular and cellular levels propagate through higher levels of brain functional organization and mediate the autism-associated phenotypes^14^.

Behavioral and cognitive phenotypes in psychiatric disorders are likely driven by the dysfunction of particular brain circuits, which consist of functionally related and anatomically connected brain structures populated by distinct neuronal cell types that mediate circuit-specific information processing. Functional brain circuits often contain multiple feed-forward and feed-back connections that facilitate efficient information processing and integration^15^. Notably, our previous analyses of neuron types impacted by ASD-associated mutations have implicated several key components of the cortical-striatal-thalamic circuits (CSTC)^13^. The CSTC loops are known to mediate diverse motor, emotional, and habit-forming behaviors that are frequently compromised in ASD^16^. The cortical-striatal circuits have been also implicated based on behavioral analysis of genetic models of highly penetrant ASD mutations in mice^17,18^. Furthermore, the disruption of neuron types facilitating the communication between cortical-subcortical and cortical-cortical regions have been associated with ASD^8,12,13^. Brain imaging and fMRI studies of ASD and other psychiatric disorders also revealed hyper– and hypo-connectivity between diverse brain regions^19,20^. In this context, to discover the brain circuits and associated cell types affected in ASD and other mental disorders, it is important to integrate, in an unbiased and genome-wide fashion, the accumulated genetic information with brain-wide single-cell and spatial transcriptome and connectome data.

To identify brain circuits and related cell types preferentially targeted by ASD mutations, we present in this paper the first approach, to our knowledge, that integrates high-resolution spatial transcriptomics with brain-wide anatomical connectome and with genome-wide genetic data; we refer to our approach as GENCIC (for Genetics and Expression iNtegration with Connectome to Infer affected Circuits). Mesoscale brain-wide connectome and expression data at high spatial resolution, and in the same coordinate system, are not currently available for humans. Fortunately, the corresponding high-quality mouse datasets have been generated by the Allen Institute^21–23^, and these valuable resources make our approach possible. Notably, multiple studies of wild-type mouse strains and mouse models of human genetic insults associated with psychiatric disorders demonstrated that many relevant brain circuits are evolutionarily and functionally conserved in mice^24,25^. In GENCIC, we first identify the mouse orthologs of the ASD genes that have been implicated in the Simons Foundation Powering Autism Research for Knowledge (SPARK) study^5^ (see Figure 1). We then use spatial transcriptomics data from mice^21^ to identify the brain structures with specific expression of ASD genes weighted by the enrichment of the disorder-associated mutations. We next use the brain-wide connectome data^21^ to search for cohesive brain circuits connecting the brain structures with strong ASD mutation biases. We investigate the connectivity and structural properties of the targeted brain circuits, and the range of interrelated roles they likely play in brain function and ASD etiology. By analyzing the latest Allen Brain Cell Atlas – a comprehensive spatial single-cell dataset containing 5,322 molecularly defined cell types across the entire mouse brain^23,26^ – we then identify strongly affected circuit cell types. Overall, our approach implicates spatially distributed circuits and related cell types in the pathophysiology of ASD and suggests why particular phenotypes may be commonly associated with the disorder.

**Figure 1:**
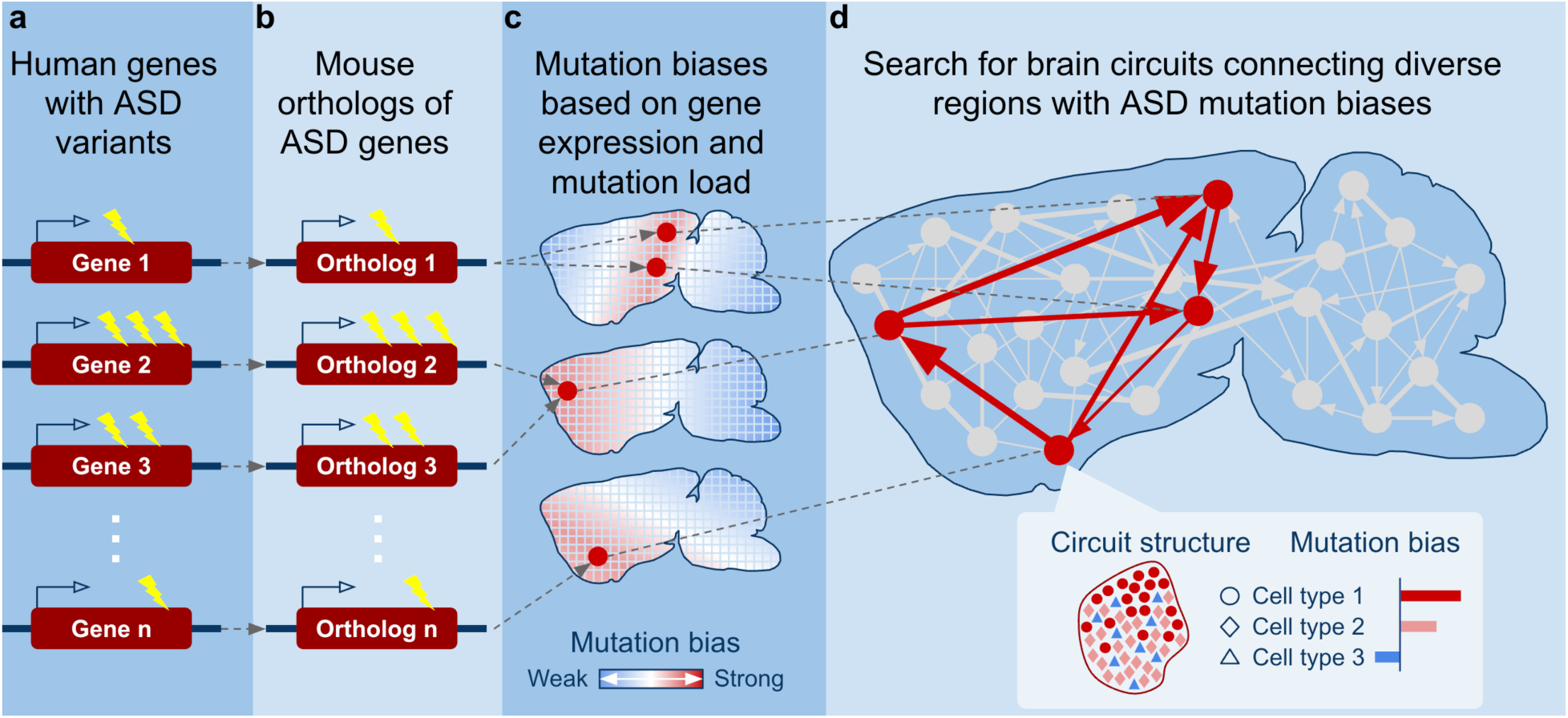
Illustration of the developed approach for Genetics and Expression iNtegration with Connectome to Infer affected Circuits (GENCIC). **a**.) High-confidence ASD genes are identified using genome-wide mutation data and their relative mutational contributions are calculated. b.) Implicated human genes are mapped to their mouse orthologs. c.) Based on mouse high-resolution spatial transcriptomics data, ASD biases are calculated for distinct anatomical brain structures. d.) Using brain-wide connectome data (gray arrows), a global optimization algorithm searches across the entire brain for cohesive functional circuits (red arrows) connecting brain structures with high ASD mutation biases. The statistical significance of the ASD circuits is estimated using circuits obtained by GENCIC based on mutated genes in unaffected siblings. Cell type composition and biases were analyzed for each circuit structure utilizing spatial single-cell data.

## Results

To identify brain circuits preferentially affected in ASD, we used genes and mutations identified in the recent meta-analysis of 42,607 ASD probands and 8,267 siblings^5^. The SPARK study prioritized 61 ASD genes with genome-wide significance based on *de novo* mutations in the affected probands. We mapped the implicated human genes to their mouse orthologs using the Mouse Genome Database^27^. Using brain-wide and genome-wide spatial transcriptomics data from the Allen Mouse Brain Atlas^21^ based on *in situ* hybridization (ISH), we then calculated the expression biases for mouse orthologs of human ASD genes across 213 neural structures spanning the entire mouse brain (Figure 1, Supplementary Figure 1). As the source of brain-wide anatomical connectome data, we utilized the Allen Mouse Brain Connectivity Atlas^22^, which was obtained using viral tracing of axonal projections. Both the connectome and transcriptome atlases were developed using the same reference model of neuroanatomical brain partitions, thus allowing us to integrate the data in GENCIC and to identify the cohesive brain circuits targeted by ASD mutations (Figure 1).

Genes associated with ASD and other psychiatric disorders are known to be highly expressed in the brain^13^, and, not surprisingly, all brain structures displayed higher expression levels of ASD-associated genes compared to genes with mutations in unaffected siblings (Wilcoxon signed-rank test P-value < 10^−10^, Supplementary Figure 2, see Methods). Therefore, to identify the brain structures and circuits that are preferentially targeted by ASD mutations, we focused on the specificity of the mutational impact. To that end, we quantified how specific is the mutation-weighted expression of ASD genes across the considered brain structures compared to the expected specificities for random genes with similar brain expression (see Methods, Supplementary Figure 1). For each ASD gene and each brain structure, we first calculated the Z-score characterizing the expression level in that structure compared to all other brain structures. The expression specificity score for each ASD gene was then calculated as the deviation between the specificity Z-score for that gene and the distribution of expected Z-scores of random genes with similar levels of overall brain expression (see Methods). The implicated ASD genes harbor various numbers of *de novo* likely gene-disrupting (LGD) and damaging missense (Dmis) mutations, and therefore have different contributions to the population risk of ASD; in addition, LGD and Dmis mutations have different average penetrances^4,5^. To account for these differences, we weighted each ASD gene by the number of observed *de novo* mutations and by the enrichment of each mutation class. The final ASD mutation bias for each brain structure was then calculated as the mutation-weighted average of the expression specificities for all implicated ASD genes (see Methods).

Overall, the ASD mutation biases were significantly stronger than biases in unaffected siblings (P-value = 2×10^−4^, Figure 2a, Supplementary Figure 3). Consistent with the broad genetic impact in ASD^13,28^, we found that the autism mutations displayed significant biases (FDR < 0.1, see Methods) to diverse brain structures from the cortex, thalamus, striatum, hippocampus, and amygdala (Supplementary Table 1). Expanding the list of ASD genes, for example using the 159 *de novo* enriched genes with genome-wide P-value < 10^−3^ from the SPARK study or the 102 genes implicated in the recent ASC consortium study^4^, resulted in very similar mutation biases (Pearson’s r = 0.97, P-value < 10^−10^, and Pearson’s r = 0.99, P-value < 10^−10^, respectively, Supplementary Figure 4 a-b). Despite a substantially higher burden of LGD and damaging *de novo* mutations in females, consistent with the female protective effect^9,29^, we observed similar brain structure biases due to mutations observed in male and female probands (Supplementary Figure 4e, Pearson’s r = 0.91, P-value < 10^−10^). Because the calculated mutation biases reflect the differences in the expression specificity between the ASD genes and expression-matched random genes, the biases were generally insensitive to the variability of neuronal composition across the brain; for example, the mutation biases remained similar when we normalized gene expression for neuronal density and neuron-to-glia ratio (Pearson’s r = 0.85, P-value < 10^−10^, and Pearson’s r = 0.88, P-value < 10^−10^, respectively, Supplementary Figure 4c-d). Overall, these results demonstrate that the genetic impact of ASD mutations is significantly more focused on a particular set of brain structures compared to unaffected siblings.

**Figure 2:**
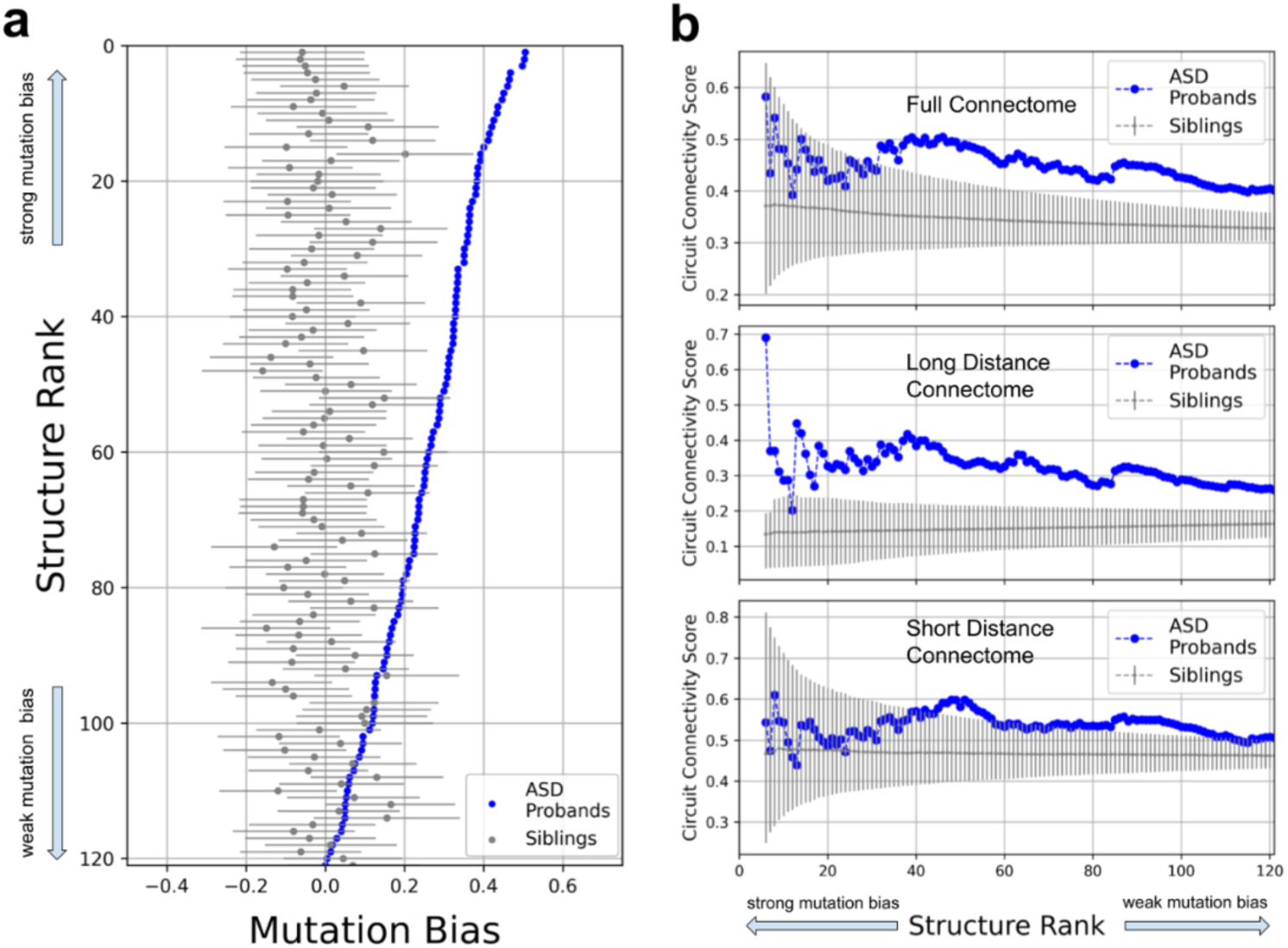
ASD mutations are significantly biased towards strongly anatomically connected brain structures. **a.)** Brain structures ranked from top to bottom (Y-axis) by the strength of their ASD mutation biases (X-axis). Each blue point corresponds to a distinct brain structure; only brain structures with positive ASD mutation biases are shown. For each structure, the corresponding mutation biases in unaffected siblings are shown in gray, with error bars representing the standard deviations across randomly subsampled sets of sibling genes (see Methods). **b.)** The structures ranked by the strength of their ASD mutation biases from left to right (X-axis). The Y-axis shows the circuit connectivity score (CCS) calculated between all brain structures to the left of a given X-axis rank. CCS for structures ranked based on the ASD mutation biases are shown in blue. The average CCS for structures ranked based on sibling mutations are shown in gray, with error bars representing the CCS standard deviations across subsampled sibling genes. The upper panel shows the CCS profile calculated using the full connectome, the middle panel shows the CCS profile calculated using connections longer than the median distance between all pairs of brain structures, and the lower panel shows the CCS profile calculated using connections shorter than the median distance between all pairs of brain structures.

We next investigated how surprising are the anatomical connections between the brain structures with strong ASD mutation biases. The probability that a pair of structures is anatomically connected strongly depends on their physical distance^22^. Thus, for a given set of brain structures, we defined a circuit connectivity score (CCS) as the average log probability of observing a particular connectivity configuration given the set of physical distances between the structures (Supplementary Figure 5, Supplementary Table 2, see Methods). Compared to structures biased by mutations observed in unaffected siblings, the structures with the strongest ASD mutation biases showed substantially higher CCS values (Figure 2b). While we observed that CCS reaches a maximum for the 46 structures with strongest ASD mutation biases (P-value = 0.03, Supplementary Figure 6a), CCS values remained significantly higher for ASD compared to unaffected siblings across a broad range of considered structure sizes (with P-value < 0.05, Supplementary Figure 6e). We also observed similar and statistically significant results when we considered an alternative circuit connectivity measure based on the number of connections between brain structures (P-value = 0.04, Supplementary Figure 6b). These results demonstrate that ASD mutations not only affect specific brain structures, but that they also impact structures that are strongly interconnected with each other. The significant anatomical connectivity between the structures targeted by ASD mutations suggests that they likely participate in a common set of neurobiological functions. Notably, CCS values were more significant when considering only long-distance anatomical connections between the affected structures (for connections longer than the median distance between all pairs of brain structures, P-value = 0.02 at the peak size 46, Figure 2c, Supplementary Figure 6c). In contrast, CCS values were only marginally significant when considering exclusively short-distance connections (P-value = 0.1, Supplementary Figure 6d). We explore this observation in more detail below.

A particular brain circuit affected by a set of mutations can be characterized by the specificity of mutational impact towards the circuit structures and by the connectivity strength between the structures. Therefore, in the GENCIC approach, we considered the combination of these two circuit characteristics using the Pareto front optimization^30^. The Pareto front describes a set of the most efficient tradeoffs between two or more objectives, in our case between the strength of the mutation bias towards a set of brain structures and the anatomical connectivity between these structures characterized by CCS. We used simulated annealing to globally search among all brain structures for the circuit with the highest CCS given a certain circuit size and a specified threshold for the average mutation bias towards the circuit structures (see Methods, Supplementary Figure 7). We determined the Pareto front by performing multiple simulated annealing optimizations with varying thresholds for the average mutation biases towards circuit structures (Figure 3a, Supplementary Table 3). In the optimizations, we used a circuit size of 46, corresponding to the size with the strongest CCS value (Figure 2b), but similar results were also obtained using other circuit sizes, for example 32, which corresponds to the number of brain structures with significantly strong mutation biases (FDR < 0.1, Supplementary Figure 8). To estimate the significance of the entire Pareto front for ASD, we calculated Pareto fronts based on structures impacted in unaffected siblings (see Methods). This analysis demonstrated the significance of the ASD circuit (Pareto front P-value = 2×10^−3^ for circuit size 46 and P-value = 6×10^−3^ for size 32) and thus confirmed that *de novo* ASD mutations specifically impact structures that are also strongly anatomically interconnected. For subsequent structural and functional analyses, we considered the Pareto front circuit with a substantial circuit connectivity score improvement and only a small reduction in the average mutation bias (red cross in Figure 3a).

**Figure 3:**
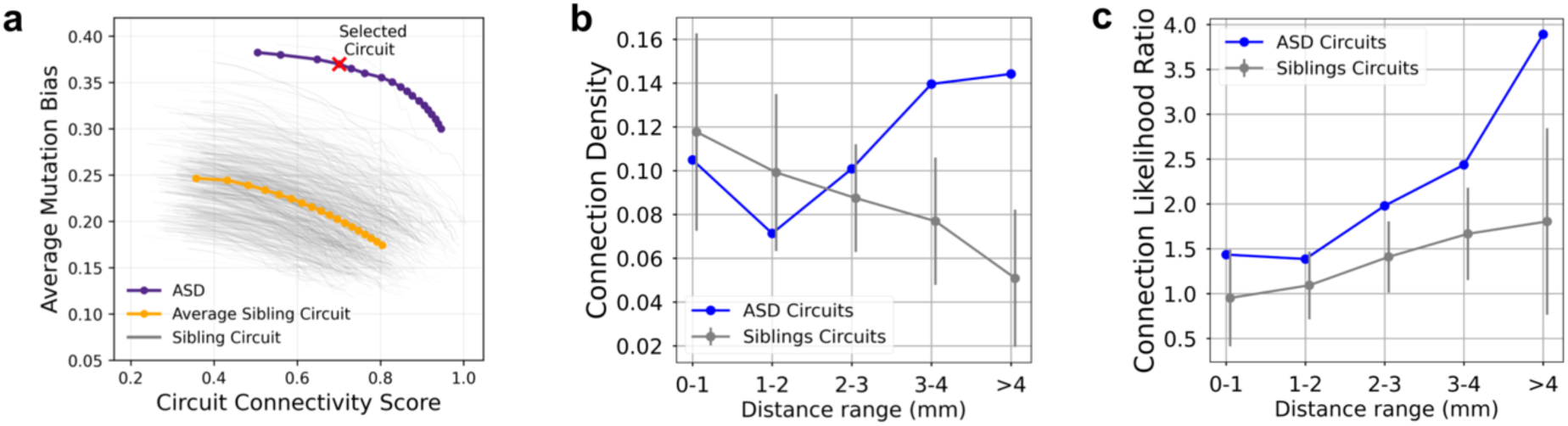
Global search identifies cohesive ASD brain circuits enriched for long-distance connections. **a**.) Pareto fronts obtained using the GENCIC algorithm. The X-axis shows the circuit connectivity scores (CCS) and the Y-axis shows the average ASD mutation bias for the brain structures forming the circuits. Simulated annealing optimizations were performed to search for the circuits with the highest CCS values while varying the minimum limit for the average mutation bias of the circuit structures. The resulting Pareto front, describing the set of most efficient tradeoffs between CCS and the ASD mutation biases, is shown in blue with points representing ASD mutation bias step sizes of 0.005; the ASD circuit selected for further analysis is marked in red. To estimate the significance of the entire Pareto front for ASD circuits (Pareto front P-value = 2×10^−3^), GENCIC searches were performed using genes with mutations in siblings. The Pareto fronts resulting from different sets of genes with mutations in siblings are shown as grey lines (see Methods), with the average sibling Pareto front shown in orange. **b.)** The densities of anatomical connections for the ASD circuit across physical distances in the mouse brain. The X-axis shows connection distances between brain structures, and the Y-axis shows the corresponding connection density, i.e., the fraction of all brain connections involved in the implicated circuits at each distance. The connection densities for the ASD circuit are shown in blue. The connection densities for circuits based on subsampled sets of sibling genes are shown in grey with error bars representing the standard deviations of densities across randomly subsampled sibling genes. The ASD circuit has similar connection density to sibling circuits at short connection distances (connection density P-value = 0.57 for distances < 3 mm), and substantially higher connection density at longer distances (P-value = 0.01 for distances > 3 mm). **c.)** The connection likelihood ratios across physical distances in the mouse brain. The X-axis shows connection distances between brain structures, and the Y-axis shows connection likelihood ratios, defined as the probability for a pair of structures in the circuit to be connected at a given distance divided by the probability for a pair of random brain structures to be connected at the same distance. The connection likelihood ratios for ASD circuits are shown in blue. The likelihood ratios for circuits based on sibling genes are shown in grey with error bars representing the standard deviations across subsampled sibling genes. Compared to circuits based on siblings, the ASD circuit has significantly higher connection likelihoods at longer distances (connection likelihood ratio P-value = 0.05 for distances > 3 mm, P-value = 0.16 for distances < 3 mm).

**Figure 4:**
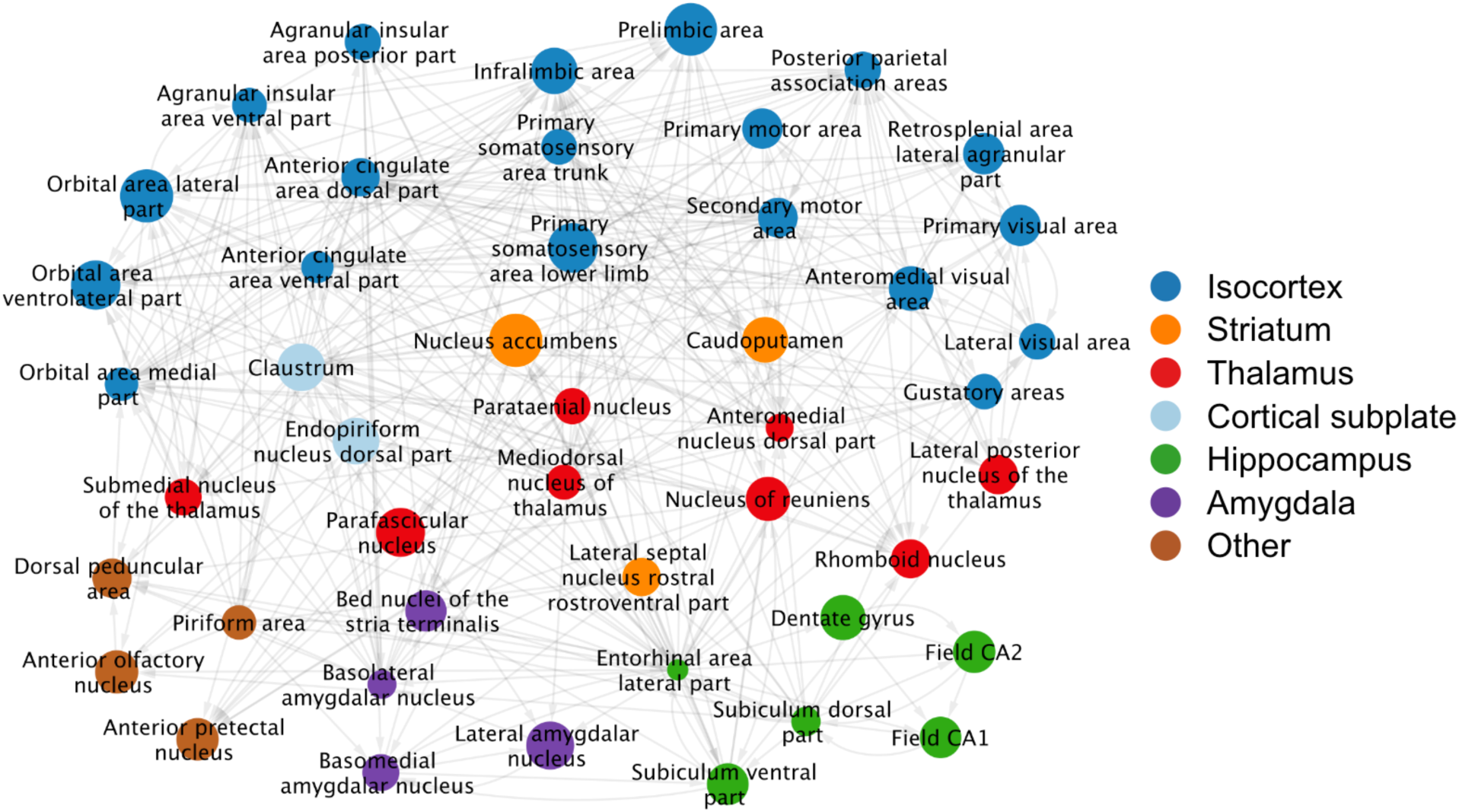
The brain structures and their anatomical interconnections forming the ASD circuit. The ASD circuit, identified using the GENCIC approach, consists of 46 distinct brain structures represented by the network nodes. Node sizes are proportional to the ASD mutation biases towards the corresponding brain structures and edge arrows indicate the direction of anatomical connections between the structures. The circuit is densely interconnected and includes ∼10% of all connectome edges. Circuit structures are colored based on broad brain regions: isocortex (dark blue), striatum (orange), thalamus (red), cortical subplate (light blue), hippocampus (green), amygdala (purple), and other brain regions (brown). Structures from the cortex and cortical subplate (dark and light blue, respectively) are primarily involved in sensory processing and decision making. The striatal and thalamic structures (orange and red) are involved in learning, reward behavior, multisensory processing and integration. The hippocampal and amygdalar structures (green and purple) provide contextual and emotional information.

Because long-distance connections strongly contribute to the significance of the ASD circuit (Figure 2b), we next explored the spatial distribution of the circuit structures and their interactions in the brain. To that end, we computed the fraction of all brain connections involved in the ASD circuit across the range of Cartesian distances (Figure 3b). This analysis showed that the fractions of short-distance (<3 mm in length) circuit connections were not significantly different (Figure 3b, connection density P-value = 0.57; see Methods) from expectation based on unaffected siblings. On the other hand, long-distance connections (>3 mm) were substantially enriched in the ASD circuit compared to unaffected siblings (connection density P-value < 0.01). To investigate whether the enrichment of long-distance circuit connections was primarily a consequence of the ASD circuit containing more distant structures, we also calculated, across distances, the likelihood ratio of connections between circuit structures (Figure 3c), which quantifies how much more likely these structures are to be connected compared to random structures at each distance (see Methods). Notably, the connection likelihood ratio of ASD structures is significantly higher at long distances (Figure 3c, connection likelihood ratio P-value < 0.05 for distances > 3mm), further supporting the functional importance of interactions between distant structures in the circuit. The circuit’s long-distance connections involve neural projections between the cortex and multiple subcortical structures, including structures in the thalamus, striatum, and amygdala, as well as distant cortico-cortical projections.

To identify circuit cell types that are strongly affected by ASD mutations, we used the latest single-cell mouse transcriptome data from the Allen Brain Cell (ABC) atlas^23^. The atlas comprises single-cell RNA-sequencing of more than 4 million cells, which were clustered into 5,322 cell types and spatially localized based on the multiplexed error-robust fluorescence *in situ* hybridization (MERFISH) analysis of an additional 4 million cells^23,26^. Notably, the ABC atlas was registered to the Allen Common Coordinate Framework (CCF), allowing us to map the distinct cell types to the corresponding circuit structures. First, to validate structural ASD biases, we reconstructed transcriptomes and mutation biases of each brain structure using single-cell ABC RNA-sequencing data (see Methods). The reconstructed mutation biases based on single-cell transcriptomics were strongly correlated with the biases based on the ISH structural data (Pearson R = 0.75, P-value < 10^−10^, Figure 5A). To identify the most strongly impacted cell types, we computed mutation biases for each of the single-cell clusters in the ABC atlas (see Methods). We identified in this analysis 206 significantly biased (FDR < 0.05) cell types (Figure 5bc, Supplementary Figure 9, Supplementary Table 3), including a heterogenous set of neuron types from multiple cortical and subcortical structures (see below). These results are consistent with and significantly expand our previous work and the work of others implicating various cell types perturbed in autism^4,8,12,13^, and suggest that the ASD mutation biases converge on a set of strongly interconnected neural structures and specific neuron types associated with these structures. To understand the set of common neurobiological functions likely underlined by the implicated circuit, we next describe the circuit structures, cell types and key interactions between them.

**Figure 5:**
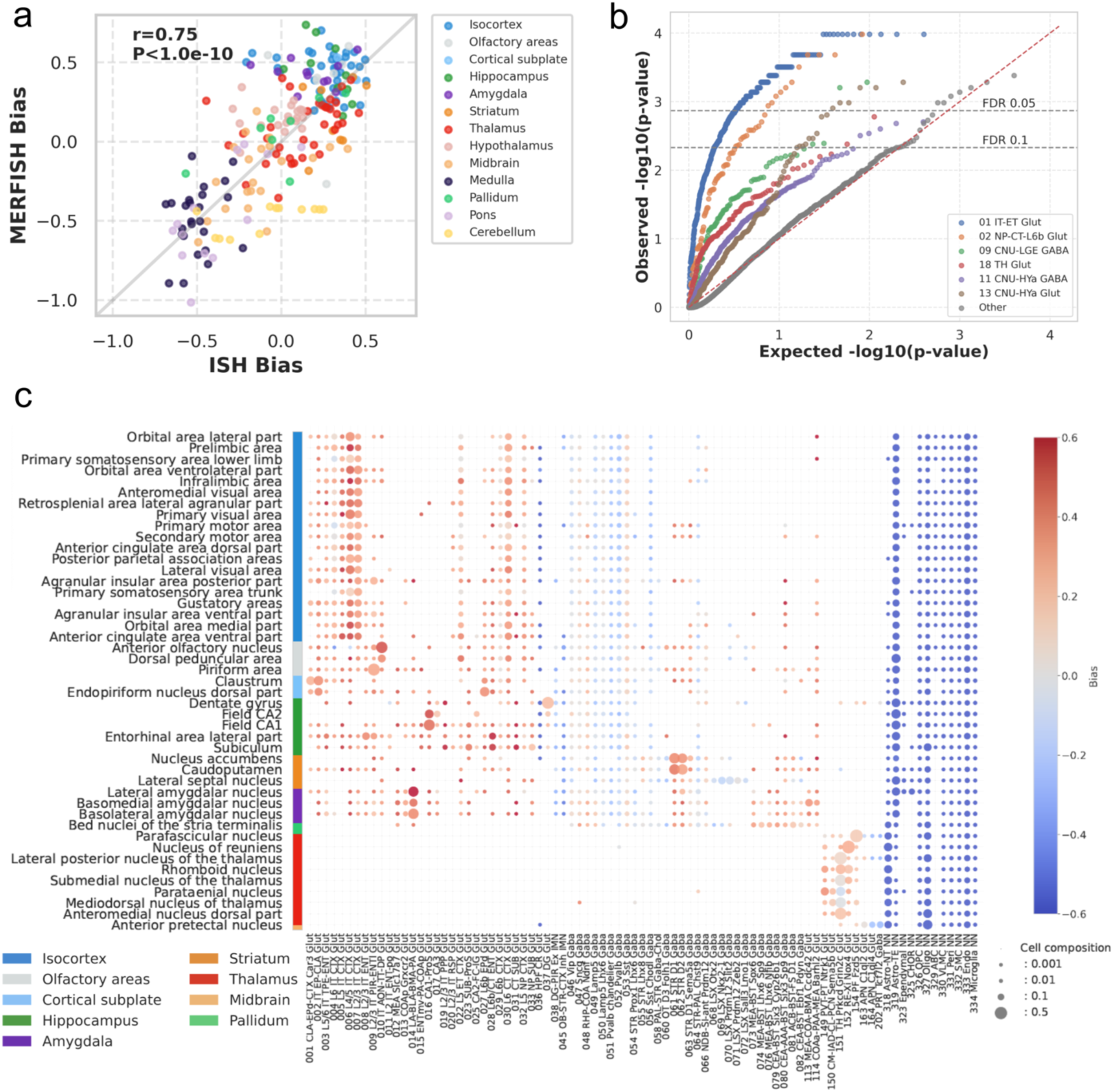
ASD mutation biases for cell types from the ABC atlas of the mouse brain. **a.)** Spearman’s correlation between brain structure ASD mutation biases calculated using ISH data (X-axis) and using aggregated ABC atlas spatial single-cell data (Y-axis). Each point represents a brain structure, colored by brain region. **b.)** QQ plot of ASD cell type mutation biases, colored by major cell type class with strong ASD biases. Each point represents a cell type cluster. The Y-axis shows the –log_10_ mutation cell type bias P-value calculated based on biases observed in unaffected siblings and the X-axis shows the expected P-value quantile. **c.)** Cell type composition and mutation biases of ASD circuit structures. Cell types with more than 500 structurally mapped cells in ASD circuit structures are shown at the ABC subclass level on the X-axis, and corresponding circuit structures are shown on the Y-axis. Each dot represents a cell type, with its size indicating the fraction of that cell type in a corresponding brain structure; the dot color represents ASD mutation bias towards the cell type. Cell types shown (X-axis) include: intratelencephalic (IT), extratelencephalic (ET), and near-projecting (NP) glutamatergic neurons (labelled 001-037); immature neurons in hippocampus and olfactory areas (038-045); medial and caudal ganglionic eminence (MGE/CGE) derived interneurons (039-053); medium spiny neurons and interneurons of the striatum (054-064); subpallial amygdala glutamatergic and GABAergic neurons (068-114); thalamic glutamatergic neurons and midbrain-derived inhibitory neurons (149-164); and non-neuronal brain cells (318-334). Complete data can be found in Supplementary Table 4.

The identified circuit includes multiple sensory areas of the cortex, including olfactory, visual, and somatosensory areas (Figure 4, dark blue). This pattern is consistent with multi-modal sensory abnormalities frequently observed in autism^31^. Strong impact is also observed in structures of the medial prefrontal cortex (mPFC), such as the prelimbic and paralimbic cortices, which are involved in decision making and regulation of reward-motivated behaviors^32^. Key positions in the circuit are occupied by several interrelated cortical areas with extensive connections to the limbic system, such as the orbital areas, anterior cingulate cortex (ACC), and insular cortex (Figure 4, dark blue). The orbitofrontal cortices encode information about rewards and emotions associated with alternative decisions and outcomes^33^. The ACC plays a crucial role in mediating the cortical control of emotions and in evaluating the salience of emotional and motivational information^34^. The insular cortex is involved in interoception and in processing basic emotions such as disgust, anger, and anxiety^35^. Another major hub of the circuit is the claustrum, which reciprocally interacts with multiple cortical areas and plays an important role in salience detection and shifting of attention^36^. Cortical excitatory neurons from both the upper and deep layers show strong ASD biases, including layer 2/3 and 4/5 intratelencephalic (IT), layer 6 cortico-thalamic, as well as near-projecting neurons (Figure 5c, Supplementary Figures 9 and 10a, Supplementary Figure 11c). While generally less impacted than excitatory neurons, various inhibitory neurons also show significant ASD mutation biases (Figure 5c, Supplementary Figure 10c). These inhibitory neurons, which originate from the medial and caudal ganglionic eminences (MGE/CGE), are usually shared across the cortex, hippocampus, and amygdala^26^, and are crucial for cognitive and emotional regulation^61–63^.

The cortical areas of the circuit communicate directly with the ventral and dorsal striatum, i.e., the nucleus accumbens (NAc) and caudoputamen (CP) (Figure 4, orange). ASD genetic insults to these areas likely contribute to deficits in social communication, restricted interests, and repetitive behaviors frequently observed in ASD^37^. Through connections with multiple cognitive and emotional cortices as well as the hippocampus and amygdala, the NAc is involved in dopamine-based reinforcement learning and in processing reward, novelty, and motivation^38^. The NAc also interacts with the lateral septum, another subcortical circuit structure involved in reward and social behavior^39^. Strong mutational impact to the NAc may contribute to the social and cognitive deficits observed in ASD. The CP strongly interacts with the motor cortical structures and regulates planning and execution of movements, processes commonly compromised in autism^40^. D1 and D2 medium spiny neurons (subclass 061 STR D1 and 062 STR D2, Figure 5c, Supplementary Figure 11a) in the NAc and CP are among the most strongly affected neuron types (Figure 5bc, Supplementary Figure 10d). These neurons jointly facilitate and modulate reward-related behaviors, and when disrupted, give rise to social deficit and repetitive behaviors^41^.

The circuit also contains several limbic and cognitive thalamic nuclei (Figure 4, red) that facilitate cortico-cortical and cortico-limbic communications^42,43^. The mediodorsal nucleus (MD) is a major component of the cortico-striatal-thalamo-cortical (CSTC) loops, which may underlie multiple neurobiological functions affected in autism, including the execution of movement, attention, behavior flexibility, and decision-making^44,45^. Among the PF and MD, glutamatergic neurons, such as Fzd5-expressing neurons, show strong ASD biases (Figure 5c, Supplementary Figure 10d, Supplementary Figure 11b). The lateral posterior nucleus interacts with visual and association cortical areas of the circuit and participates in multisensory processing and detection of visual salience^46^. The anteromedial thalamic nucleus interacts with the mPFC to regulate motivation and goal-directed behaviors^47^. The reuniens (RE) and rhomboid nuclei (RH) play important roles in gating the flow of information between multiple cortical and hippocampal areas of the circuit^48^. Among RE and RH nuclei, several glutamatergic neurons, such as Nox4– and Ntrk1-expressing, show strong ASD mutation biases (Figure 5c, Supplementary Figure 10d).

The implicated circuit likely integrates into decision making relevant contextual information, including memory and emotional engagement. ASD probands often have difficulty understanding emotional context^49^ and exhibit various memory problems^50^. Processing and integration of contextual information is supported by the hippocampal (Figure 4, green) and amygdalar nuclei of the circuit (Figure 4, purple), which interact with each other^51^ and with multiple other circuit structures. The CA fields of the hippocampus play key roles in facilitating learning and memory, and the subiculum mediates the flow of information from the hippocampus^52^. Numerous glutamatergic neurons that are either enriched in the hippocampus or are broadly expressed throughout the limbic system are strongly affected by the ASD mutation biases (Figure 5c, Supplementary Figure 10ad, Supplementary Figure 11b). The basolateral amygdala (BLA) is involved in processing and formation of emotional memories^53^, and the basomedial amygdala (BMA), together with the bed nucleus of the stria terminalis (BST)^54^, participates in processing fear and anxiety^55^. Multiple glutamatergic neurons of the amygdala show strong ASD mutation biases (Figure 5c, Supplementary Figure 10ab). Various inhibitory neurons of the amygdala are also strongly biased, including D1/D2 medium spiny neurons, which play a role in reward learning and decision making^56^.

An established hallmark of ASD is the broad phenotypic heterogeneity among affected individuals^1,2,57^, with cognitive and intellectual disabilities being common comorbidities^58,59^. To understand how mutation biases towards different circuit structures relate to cognitive phenotypes, we analyzed ASD mutations in probands stratified by their full-scale intelligence quotient (IQ) scores. To that end, we partitioned probands into lower IQ (≤70) and higher IQ (>70) cohorts and calculated in each of the cohorts the ASD mutation biases for all circuit structures (Figure 6). Interestingly, mutations in the lower IQ cohort showed consistently stronger biases across all circuit structures compared to mutation in the higher IQ cohort (Wilcoxon signed-rank test, P < 10^−10^, for all circuit structures). This suggests that mutations leading to severe cognitive impairments usually have more pervasive effects on the entire circuit. The most pronounced differences between the higher and lower IQ cohorts were observed in structures from the isocortex, hippocampus, and amygdala – regions crucial for cognitive functions^60^ (Figure 6, Supplementary Table 5). These results are consistent with recent fMRI studies of ASD probands^20^ and complement our previous findings quantifying the effects of gene dosage changes on cognitive autism phenotypes^61^.

**Figure 6:**
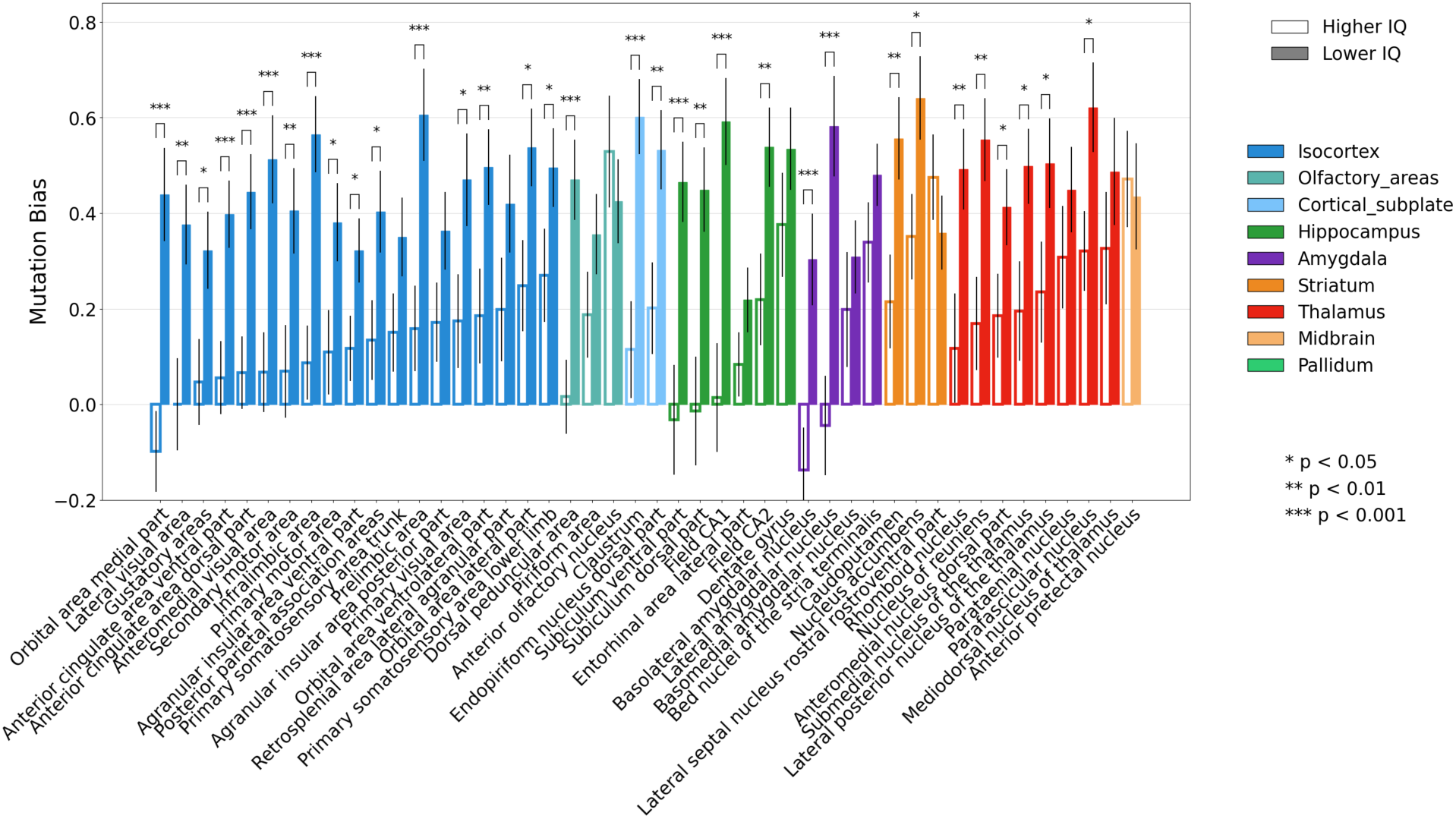
Sensitivity of the Full-Scale IQ score to ASD mutation biases across circuit structures. Comparison of the ASD mutation biases between the higher FSIQ (FSIQ>70, open bars) and the lower FSIQ (FSIQ<70, solid bars) proband cohorts across brain structures of the ASD circuit. Brain structures are color-coded by anatomical regions (isocortex, olfactory areas, cortical subplate, hippocampus, amygdala, striatum, thalamus, midbrain, and pallidum). Statistical significance of the differences between the higher and lower IQ cohorts is shown (*p < 0.05, **p < 0.01, ***p < 0.001, IQ permutation test). Error bars represent the standard error of the mean derived from a bootstrap analysis of ASD mutations (see Methods).

## Discussion

Previous functional analyses of genetic mutations associated with ASD revealed wide-ranging perturbations of genes, molecular functions, and cell types^6–13^. The results described in this paper show that, despite a widespread genetic impact, the brain structures most strongly and specifically affected in autism tend to preferentially interact with each other. These interactions are especially enriched in long-distance anatomical connections that bind the spatially distributed structures into a set of interrelated functional circuits. Moreover, we observed that the diversity of the impacted brain structures is matched by the diversity of neuron types targeted by autism mutations (Figure 5c, Supplementary Figure 11). Our analysis indicates that multiple strongly affected neuronal cell types, including layer 2/3 and 4/5 excitatory intratelencephalic, medium spiny neurons, and inhibitory interneurons, are often shared between the affected circuit structures. Impacting interconnected circuit structures and neuron types parallels the pattern seen at a different level of biological organization, where disease mutations target functionally connected protein and cellular networks^6,13,62^. The implicated ASD circuit (Figure 4) includes multiple cortical structures involved in cognitive and emotional control, as well as striatal and thalamic areas dedicated to information integration, learning, and decision making, and amygdalar and hippocampal regions that provide contextual and emotional information (Figure 7). Consequently, these results demonstrate that diverse disruptions in sensory integration, information processing, and decision-making lie at the core of the behavioral and cognitive phenotypes commonly observed in ASD.

**Figure 7:**
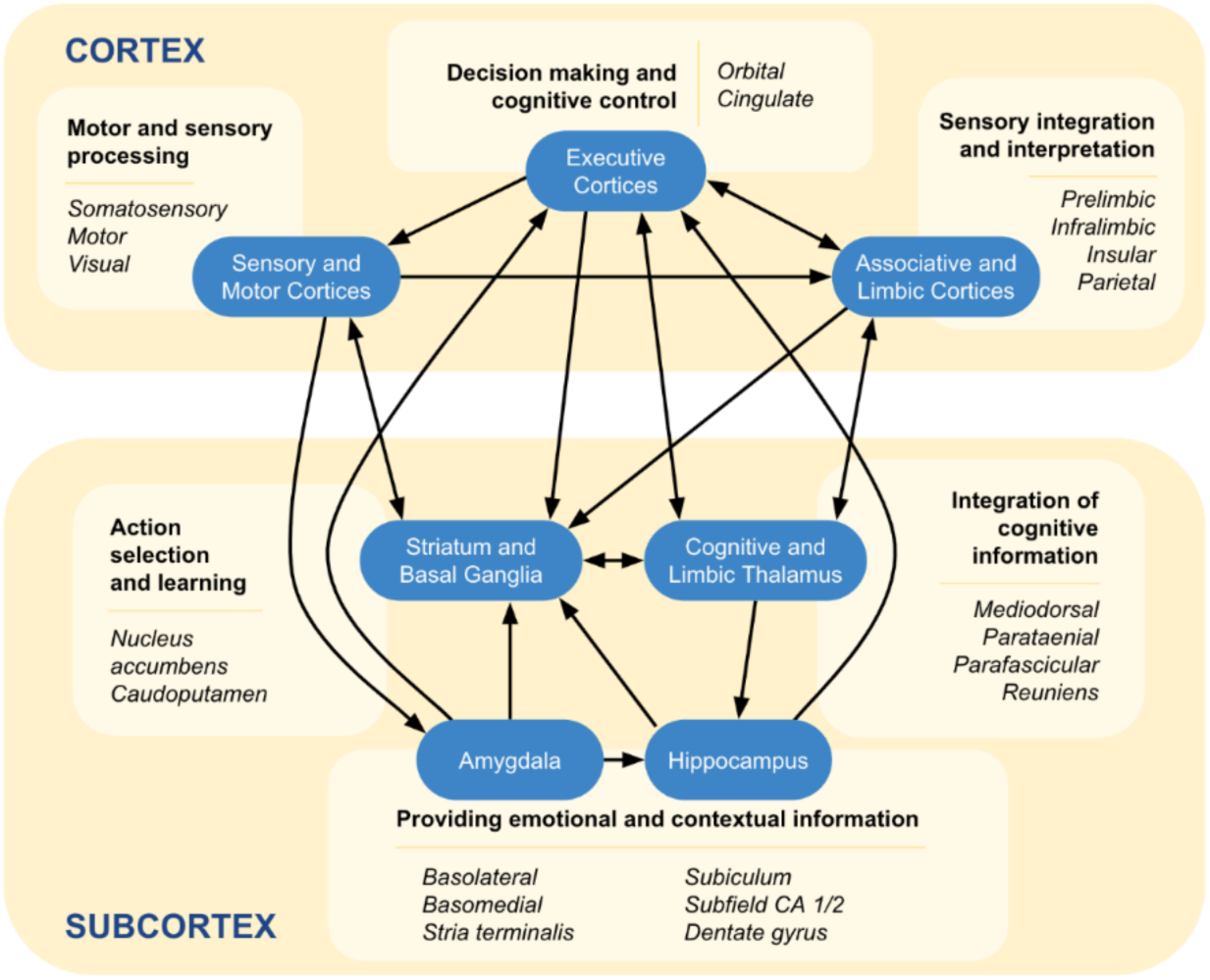
Global view of the key functional interactions between regions of the implicated ASD circuit. The ASD circuit includes multiple cortical (top yellow) and subcortical (bottom yellow) areas. The main brain areas involved in the circuit are shown in blue, and black arrows indicate the major directions of the information flow between the areas. The cortical regions of the circuit include sensory, motor, executive, associative, and limbic cortices. These regions play essential roles in information processing, sensory integration, and decision making. The subcortical circuit areas include the striatum, thalamus, amygdala, and hippocampus, which participate in information integration, action selection, and learning. Key brain structures with strong ASD mutation biases are shown in italics for each circuit brain area.

Beyond identifying ASD-specific circuits and associated cell types, we hope that this work will advance a unified circuit-based perspective on psychiatric disorders^63^, underscoring the role that circuit perturbations play in common psychiatric phenotypes^64^. By integrating genome-wide genetic data with a comprehensive spatial brain-wide transcriptome and anatomical connectome, the GENCIC approach enables unbiased discovery of circuits directly disrupted by causal mutations. This approach also complements brain imaging studies that reveal the functional changes in affected circuits^20^, as well as single-cell transcriptomics analyses that identify perturbed neuronal populations^65,66^. Because psychiatric disorders often share similar phenotypes^67^, it is likely that overlapping circuits and cell types are affected across multiple mental conditions. Moreover, as our analysis of cognitive phenotypes demonstrates, stronger impacts on specific circuit components may bias disorders toward different clinical outcomes^68^. Alongside genetic factors, environmental disturbances can be crucial in the onset and progression of psychiatric disorders, including autism, potentially converging on the same circuits. For example, social interactions can modulate hormone levels that mediate the development and function of brain circuits^69^. Notably, we found significant spatial overlap (P-value = 1.6×10^−2^, Supplementary Figure 12) between the ASD circuit structures and brain regions exhibiting high expression of the oxytocin receptor, a hormone playing a major role in mammalian social development^70^.

Finally, the circuit-level perspective is important not just for understanding the etiology of mental disorders but also for designing effective medical interventions^63^. Functions of brain circuits can be affected by multiple existing pharmaceuticals^71^. Likewise, by leveraging inherent brain plasticity, cognitive behavioral therapies can engage and modify the same circuits^72^, while circuit-targeted neurostimulation approaches, such as transcranial magnetic stimulation and deep brain stimulation, can be designed to modulate circuits’ activity and connectivity^63^. All these approaches may be synergistically applied to alleviate patient-specific disturbances within brain circuits across various psychiatric disorders.

## SUPPLEMENTARY FIGURES

**Supplementary Figure 1:**
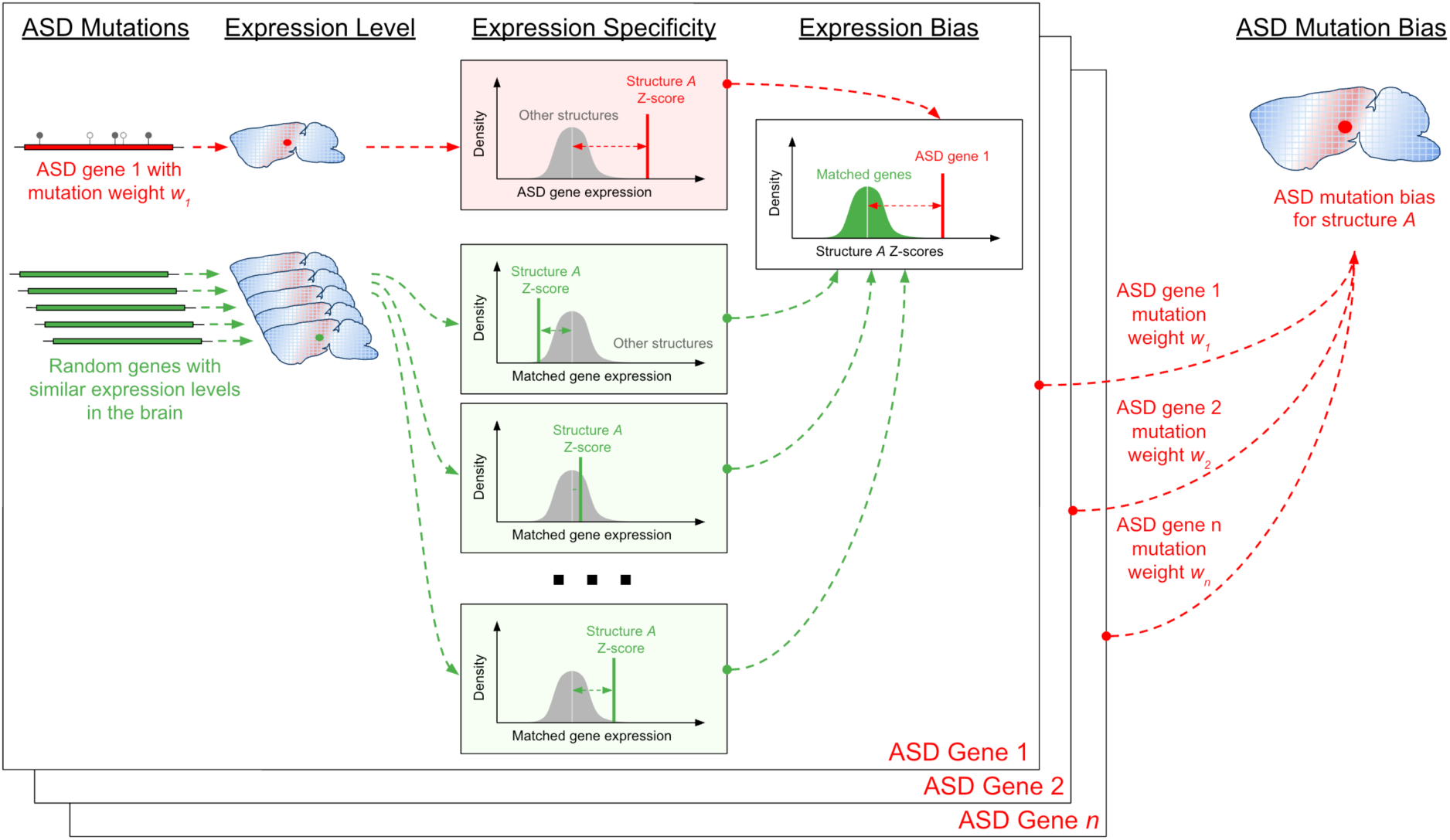
Illustration of ASD mutation bias analysis. Illustration of the gene expression analysis used to determine ASD mutation biases for each brain structure. To calculate the biases, gene expression specificities were first calculated for each gene and for each structure in the brain; the expression specificity of a gene in a brain structure was calculated as the Z-score between the gene expression level in the structure compared to the distribution of the gene expression levels in all brain structures. The specificities were calculated for each ASD gene (red) and for randomly matched genes (green) with similar expression levels in the brain. For each brain structure, expression biases were then calculated for each autism gene as the Z-score between the expression specificity of that gene and the distribution of expression specificities of random genes with similar expression level in the brain. Based on gene expression biases, ASD mutation biases were determined for each structure in the brain by computing a weighted average of mutation biases across all ASD genes; ASD gene weights were based on both the mutation class enrichments and the number of ASD mutations in each gene (see Methods).

**Supplementary Figure 2:**
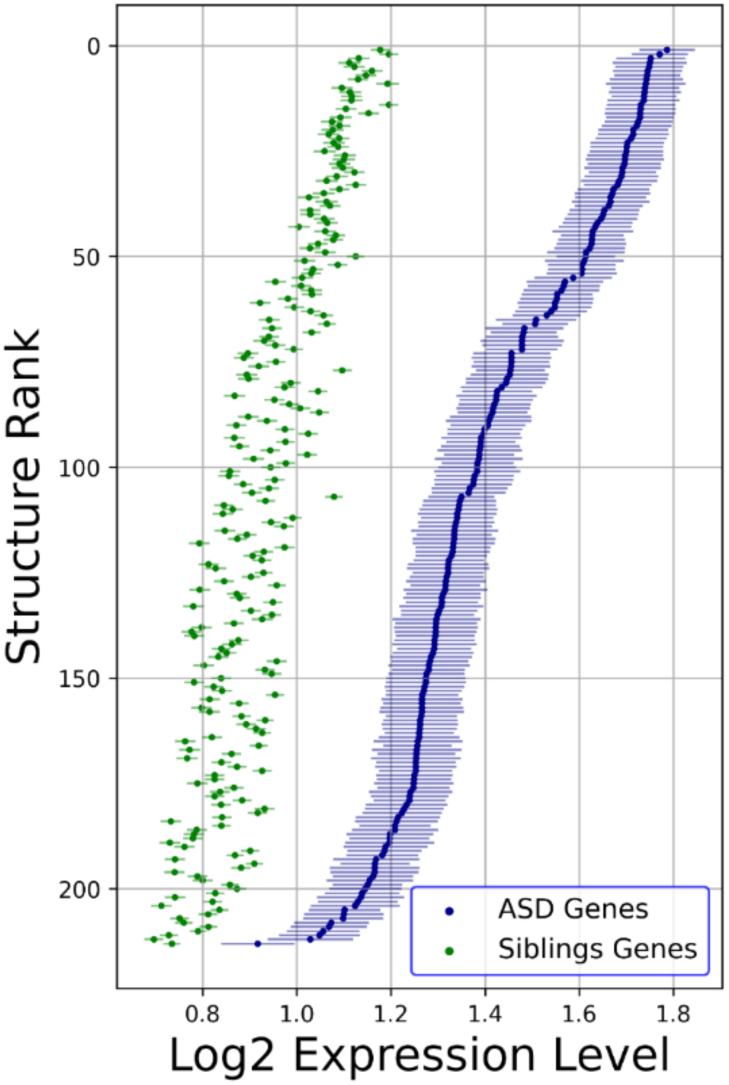
Expression levels of ASD and sibling genes across the brain. The X-axis shows the average log_2_ gene expression level in each brain structure. Each point represents a mouse brain structure, and the structures are ranked by the average ASD gene expression level from high to low along the Y-axis. Average ASD gene expression level in different brain structures are shown in blue, and average gene expression level for unaffected siblings in the same structures are shown in green. Error bars show the standard error of the average gene expression for ASD and unaffected siblings.

**Supplementary Figure 3:**
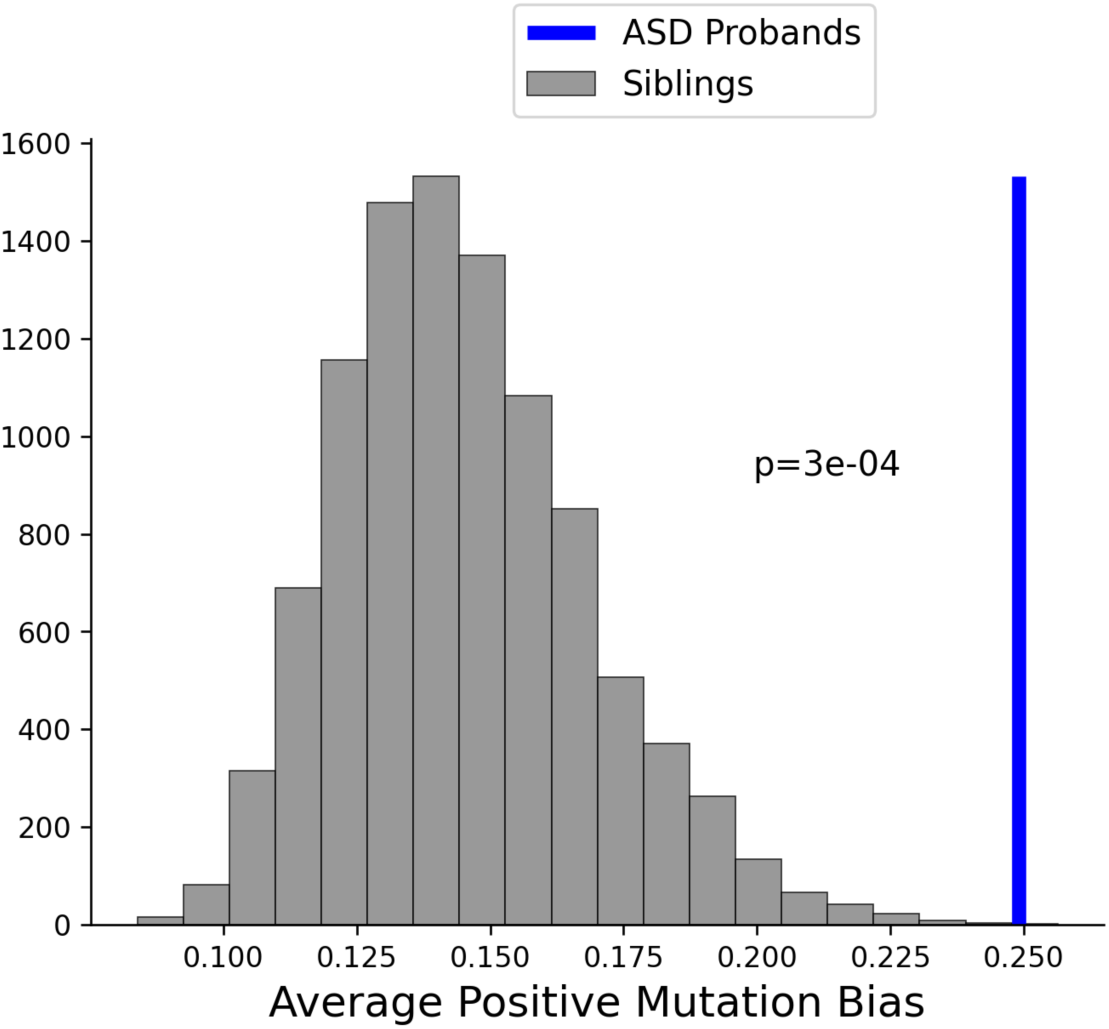
Significance of ASD mutation biases compared to mutation biases in unaffected siblings. The average ASD mutation bias in brain structures with positive biases for autism probands is represented by the vertical blue line. The distribution of mutations biases for siblings is shown in grey. To calculate the distribution of mutation biases for siblings, the genes with mutations in unaffected siblings were randomly subsampled to match the number of genes in the ASD gene set (60); The distribution of sibling mutation biases was then calculated based on 10,000 randomly subsampled sets of sibling genes (see Methods).

**Supplementary Figure 4:**
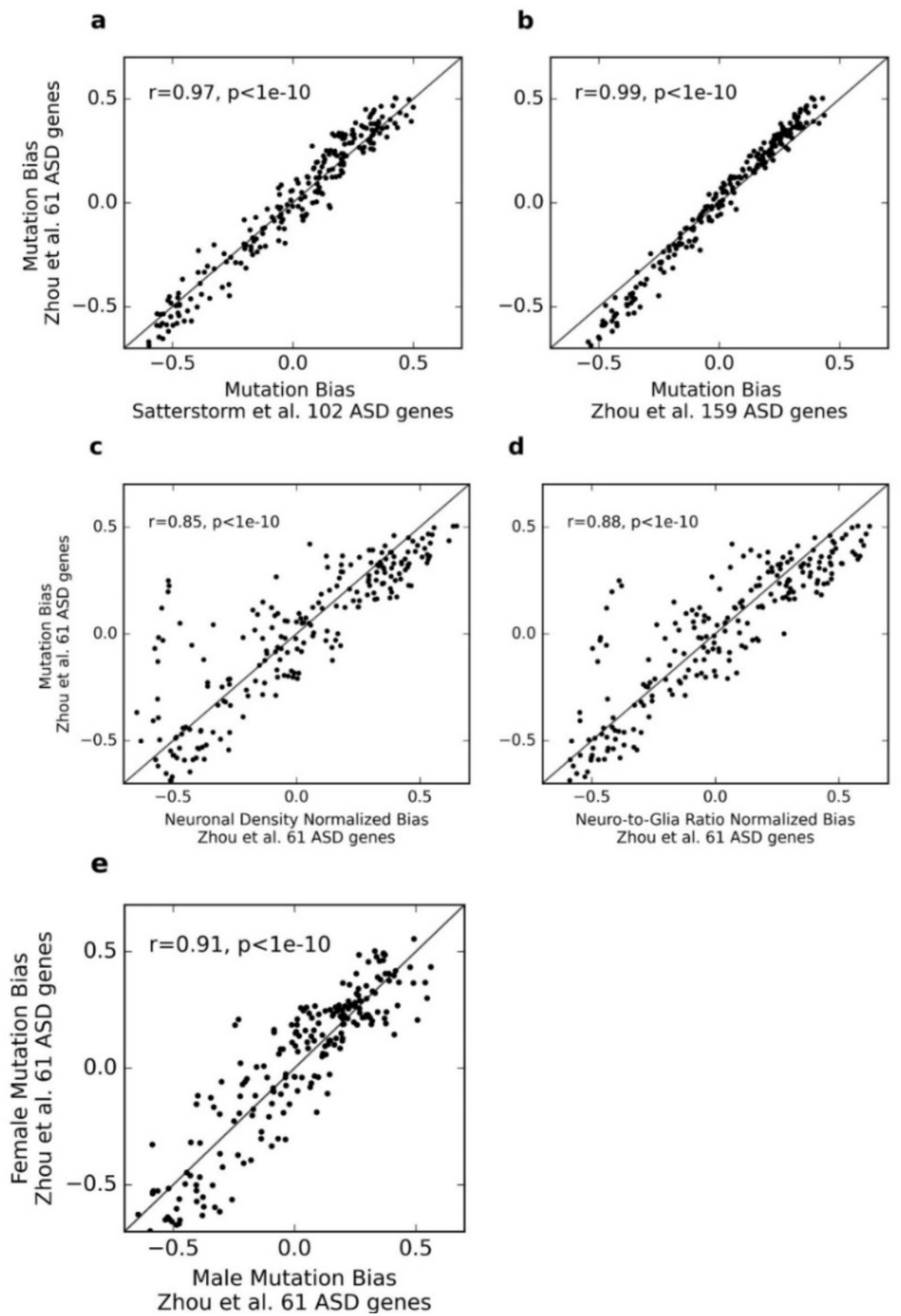
Pearson’s correlation between brain structure ASD mutation biases calculated using various datasets. Scatterplots show the correlation of mutation biases between various datasets: **a.)** SPARK DeNovoWEST exome-wide significant genes from Zhou *et al*.^5^ and ASC TADA genes with FDR<0.1 from Satterstrom *et al*.^4^, **b.)** SPARK DeNovoWEST exome-wide significant genes from Zhou *et al*. and 159 SPARK *de novo* enriched genes from Zhou *et al*., **c.)** ASD mutations from Zhou *et al*. with and without the normalization of gene expression for brain structure neuronal density, **d.)** ASD mutations from Zhou *et al*. with and without the normalization of gene expression for brain structure neuron-to-glia ratio, and **e.)** ASD mutation biases observed in female and male probands from Zhou *et al*. Diagonal lines indicate the identity relationship, X=Y. Pearson’s correlation coefficients and the corresponding P-values are shown for each correlation.

**Supplementary Figure 5:**
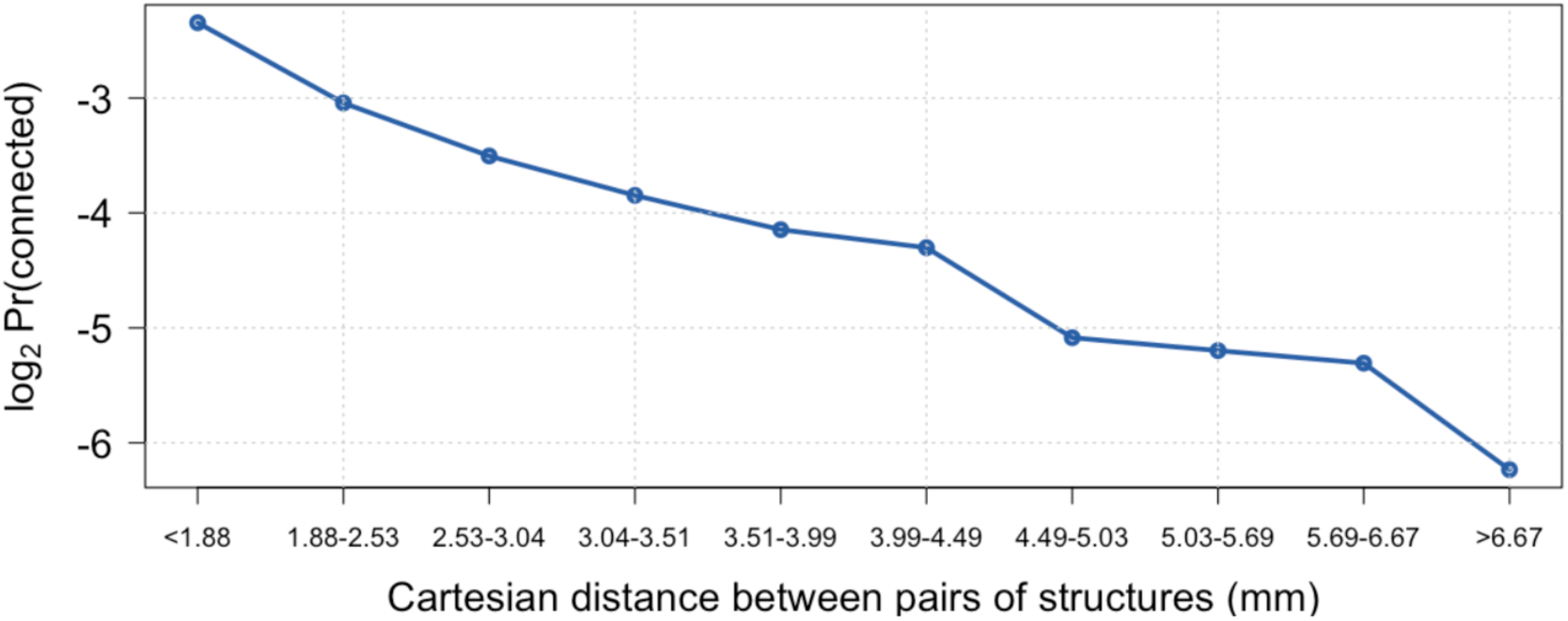
Probability of anatomical mesoscale connections between brain structures at various Cartesian distances in the mouse brain. The connection probability between structures (Y-axis) was calculated by stratifying the mesoscale connectome into 10 distance bins (X-axis) so that each bin contains the same number of brain structure pairs. The connection probability was then calculated for each bin as the fraction of connected structure pairs.

**Supplementary Figure 6:**
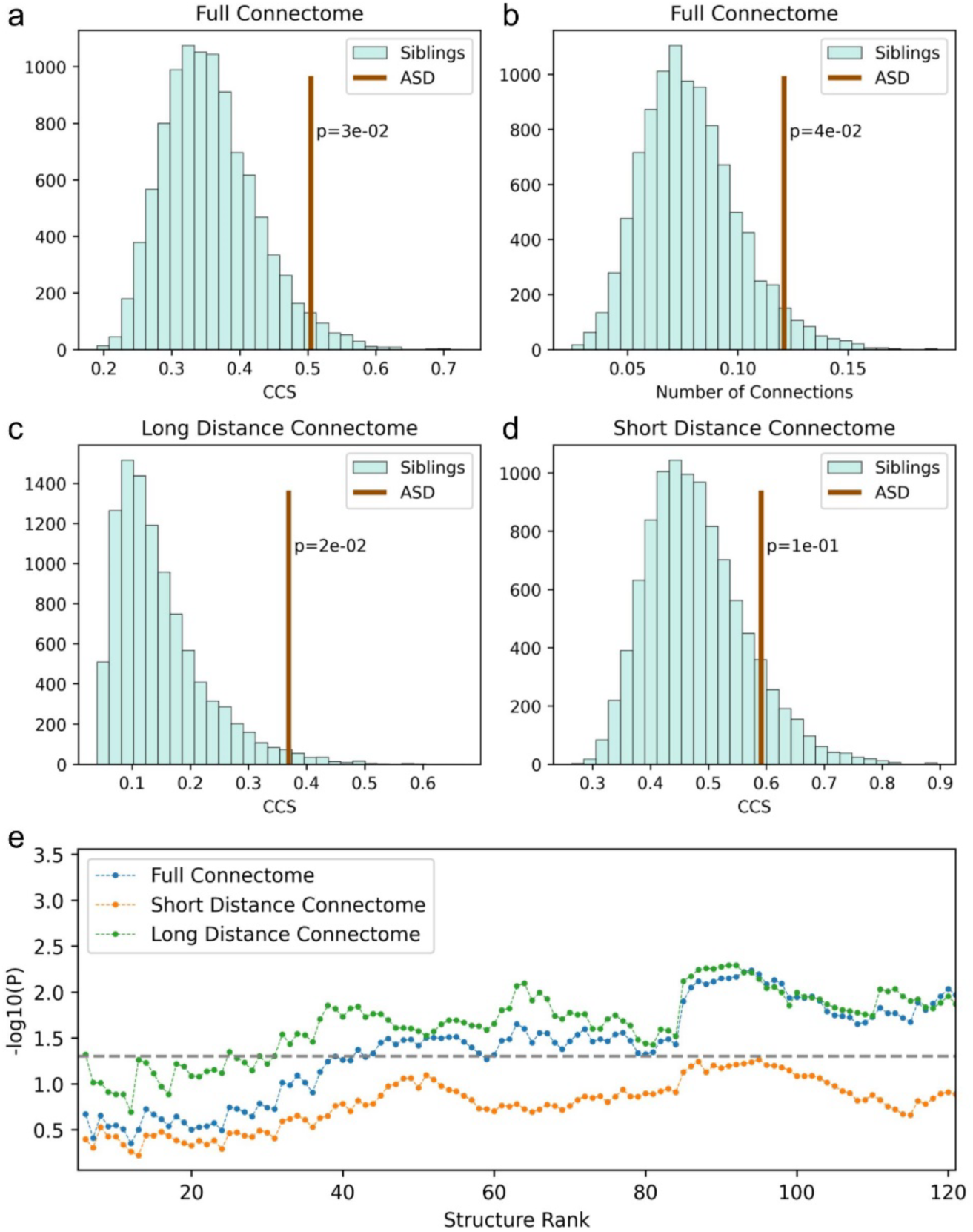
Significance of ASD circuit connectivity score (CCS) and the number of circuit connections. **a.)** CCS for the 46 structures with the strongest ASD mutation biases is represented by the vertical brown line. The distribution of CCS for siblings is shown in cyan. To calculate the distribution of CCS for siblings, the genes with mutations in unaffected siblings were randomly subsampled to match the number of genes in the ASD gene set. The distribution of sibling CCS was then calculated using the 46 structures with the strongest mutation biases for each of the 10,000 randomly subsampled sets of sibling genes. **b.)** The number of anatomical connections between the 46 structures with the strongest ASD and sibling mutation biases is shown instead of CCS. **c.)** CCS for the 46 structures with the strongest ASD and sibling mutation biases when considering only the connections spanning distances longer than the median distance between all brain structures. **d.)** CCS for the 46 structures with the strongest ASD and sibling mutation biases when considering only the connections spanning distances shorter than the median distance between all brain structures. **e.)** ASD CCS P-values were calculated for a range of brain structures with the strongest positive mutation biases. The structures were ranked based on ASD mutation biases, with the strongest biases on the left and the weakest on the right, and the significance was then evaluated for the most biased structures while incrementally varying the number of considered structures. Y-axis represents –log_10_ P-value. ASD CCS P-values calculated using the full connectome are shown in blue, using only connections spanning distances shorter than the median distance between all brain structures in orange, and using only connections spanning distance longer than the median distance between all brain structures in green. Grey dashed horizontal line indicates P-value = 0.05.

**Supplementary Figure 7:**
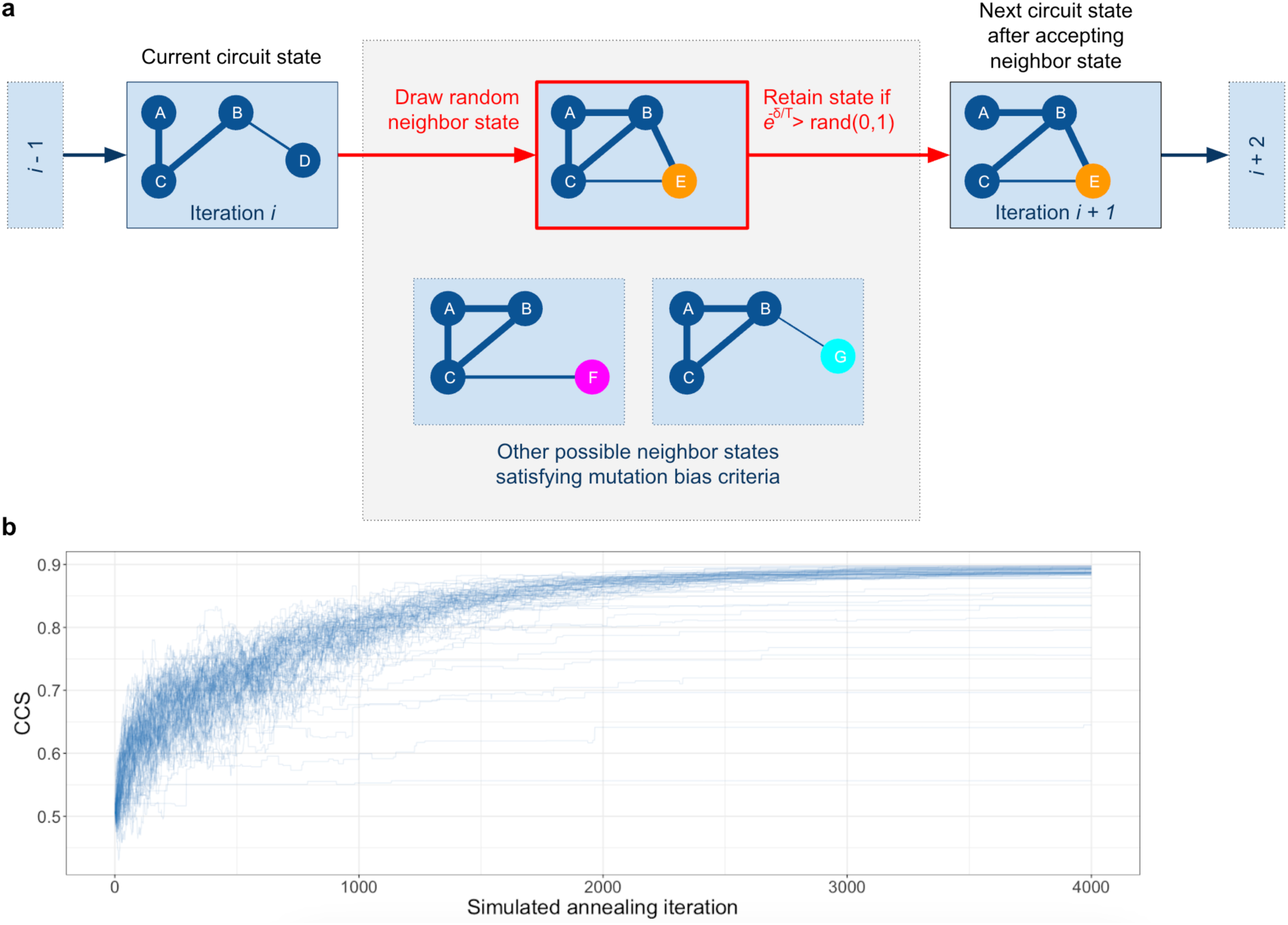
GENCIC search algorithm for cohesive brain circuits based on simulated annealing. **a.)** Illustration of GENCIC search for circuits with the maximum circuit connectivity score (CCS). The simulated annealing algorithm performed a global stochastic search by iteratively selecting a random neighbor state, i.e., attempting to replace a randomly selected circuit structure with a randomly selected brain structure not in the circuit. The proposed new state was then probabilistically accepted or rejected based on the current annealing temperature and the CCS difference between the proposed and the current state. During the annealing search, proposed new states that lowered the average mutation bias toward the circuit structures were rejected unless the average bias surpassed the minimum mutation bias threshold. **b.)** Global optimization of the CCS (Y-axis) is shown as a sequence of simulated annealing steps (X-axis) with each line representing an independent annealing search run started from different initial circuit configurations composed of randomly chosen brain structures.

**Supplementary Figure 8:**
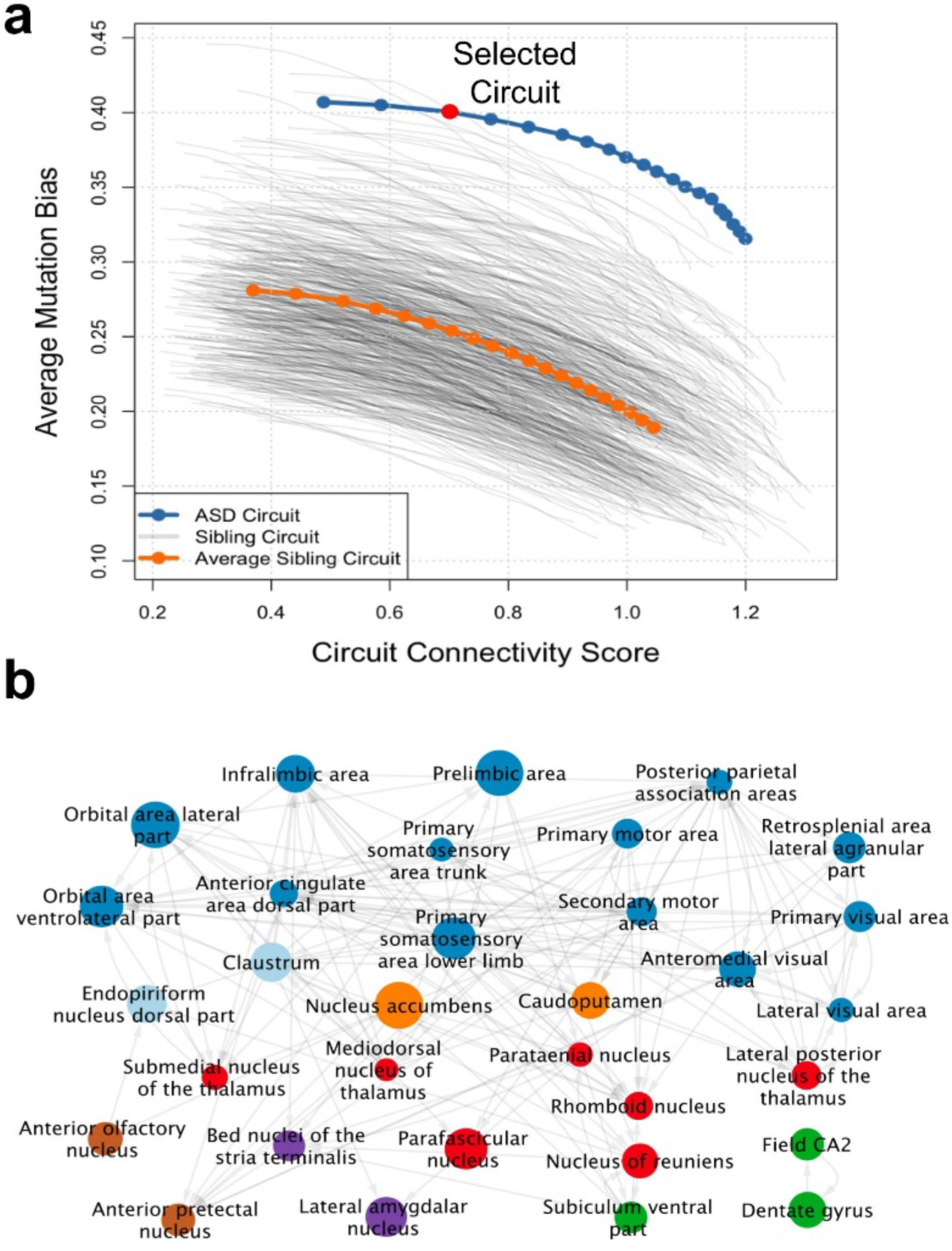
ASD circuit consisting of 32 brain structures. **a.)** Pareto fronts obtained using the GENCIC algorithm for circuits with 32 structures, corresponding to the number of structures with ASD mutation bias FDR < 0.1. The X-axis shows the circuit connectivity scores (CCS) and the Y-axis shows the average ASD mutation biases of the brain structures forming the circuits. Simulated annealing optimizations were performed to search for the circuits with the highest CCS values for different minimum values of the average mutation bias of the circuit structures. The resulting Pareto front, describing the set of most efficient tradeoffs between CCS and ASD mutation biases, is shown in blue with points representing the minimum average ASD mutation bias step sizes of 0.005. The ASD circuit shown is marked by red. To estimate the significance of the entire Pareto front for ASD circuits (P-value = 6×10^−3^), GENCIC searches were also performed using genes with mutations in unaffected siblings. The Pareto fronts for circuits resulting from subsampled sets of sibling genes are shown as grey lines (see Methods), with the average sibling Pareto front shown in orange. **b.)** The selected ASD circuit including 32 structures (nodes in the network) from the isocortex (dark blue), striatum (orange), thalamus (red), cortical subplate (light blue), hippocampus (green), amygdala (purple), and other brain regions (brown). Node sizes are proportional to ASD mutation biases of the corresponding brain structures and edges indicate the directions of anatomical connectome projections between the circuit structures.

**Supplementary Figure 9:**
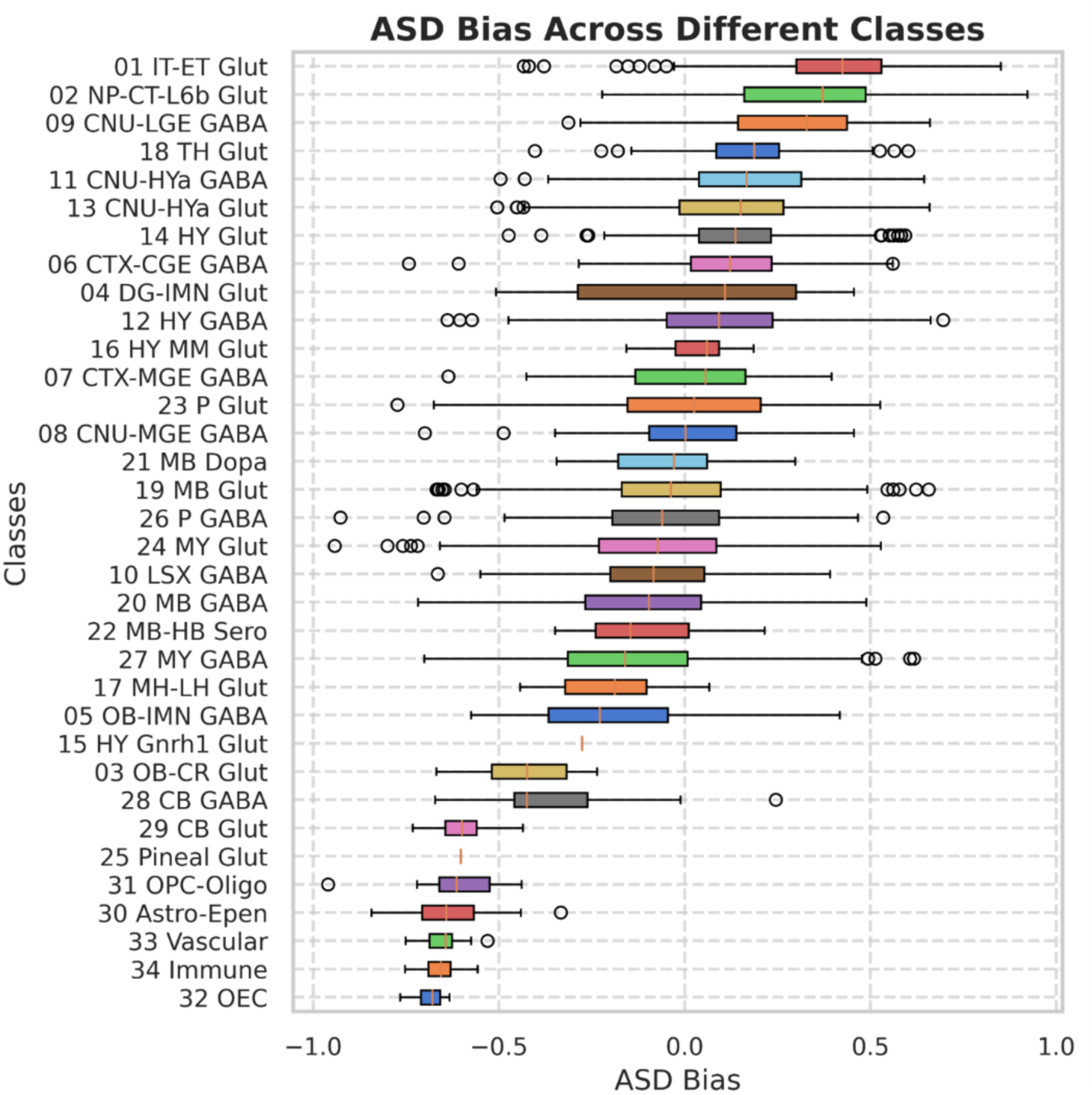
Cell type ASD mutation biases at the ABC atlas class level. ASD mutation biases for mouse brain cell types from the Allen Brain Cell (ABC) atlas grouped by cell type class. Each horizontal bar represents the distribution of mutation biases for mouse ABC cell type clusters forming a corresponding cell type class (labeled on the left). The X-axis shows the magnitude of ASD mutation biases, the Y-axis represents the 32 cell type classes in the ABC atlas. Cell type classes were sorted from top to bottom based on their median ASD mutation bias magnitude.

**Supplementary Figure 10:**
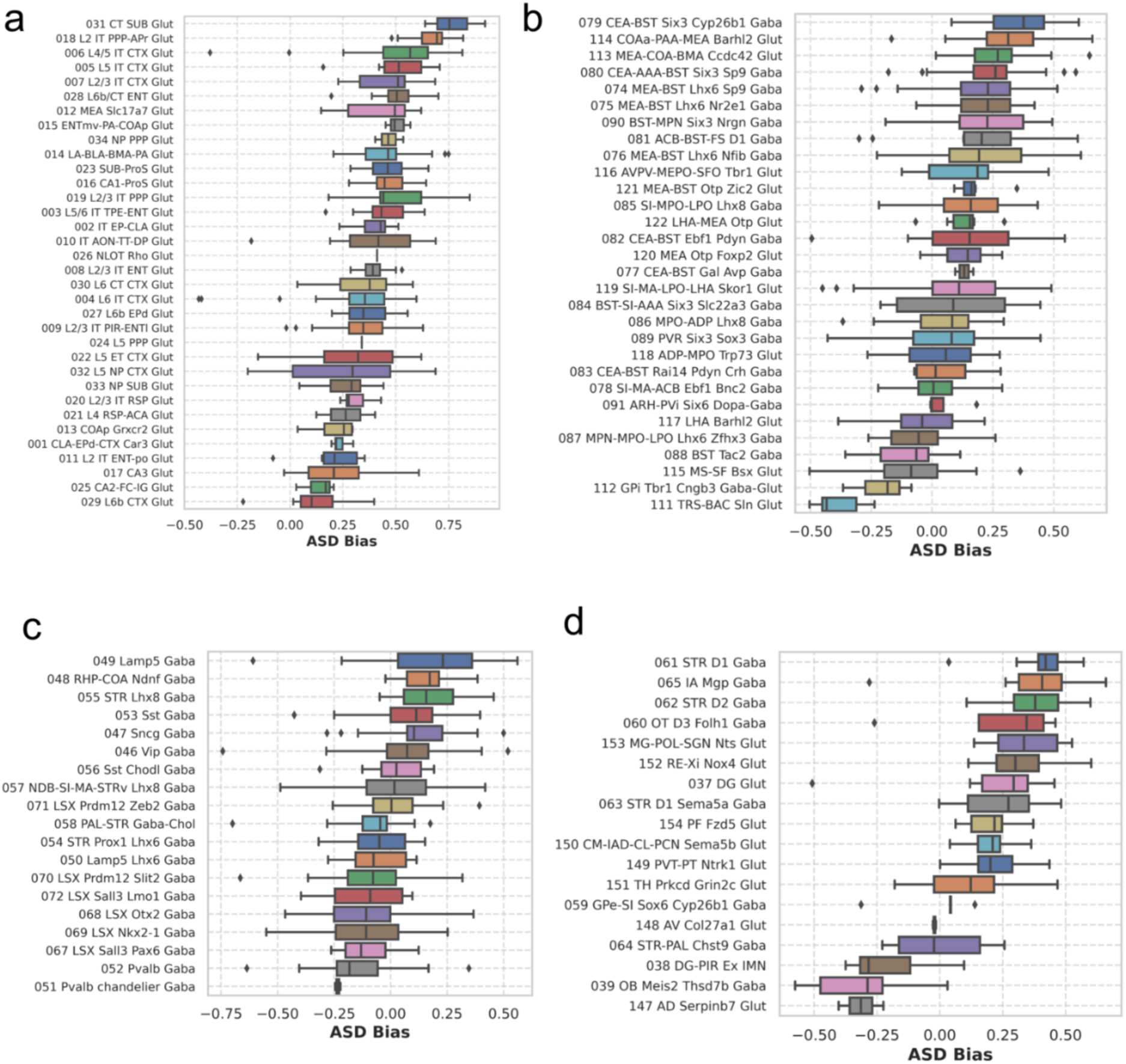
Cell type ASD mutation biases at the ABC atlas subclass level. Distribution of ASD mutation biases across neuronal subclasses in ASD-associated circuits. Each horizontal bar represents the distribution of mutation biases for the mouse brain ABC cell clusters forming a corresponding ABC cell type subclass (labeled on the left). Cell type subclasses were sorted from top to bottom based on their median ASD mutation bias magnitude. The X-axis shows the magnitude of ASD mutation biases, the Y-axis represents the ABC cell type subclasses. **a.)** ASD mutation biases for the IT-ET Glut (ABC class 01) and NP-CT-L6b Glut subclasses (class 02). **b.)** ASD biases for the CNU-HTa Glut (class 13) and GABA (class 11) subclasses. **c.)** ASD biases for the CTX-CGE GABA (class 06), CTX-MGE GABA (class 07) and CNU MGE GABA (class 08) subclasses. **d.)** ASD biases for the CNU-LGE GABA (class 09), TH Glut (class 18) and DG-IMN Glut (class 04) subclasses.

**Supplementary Figure 11:**
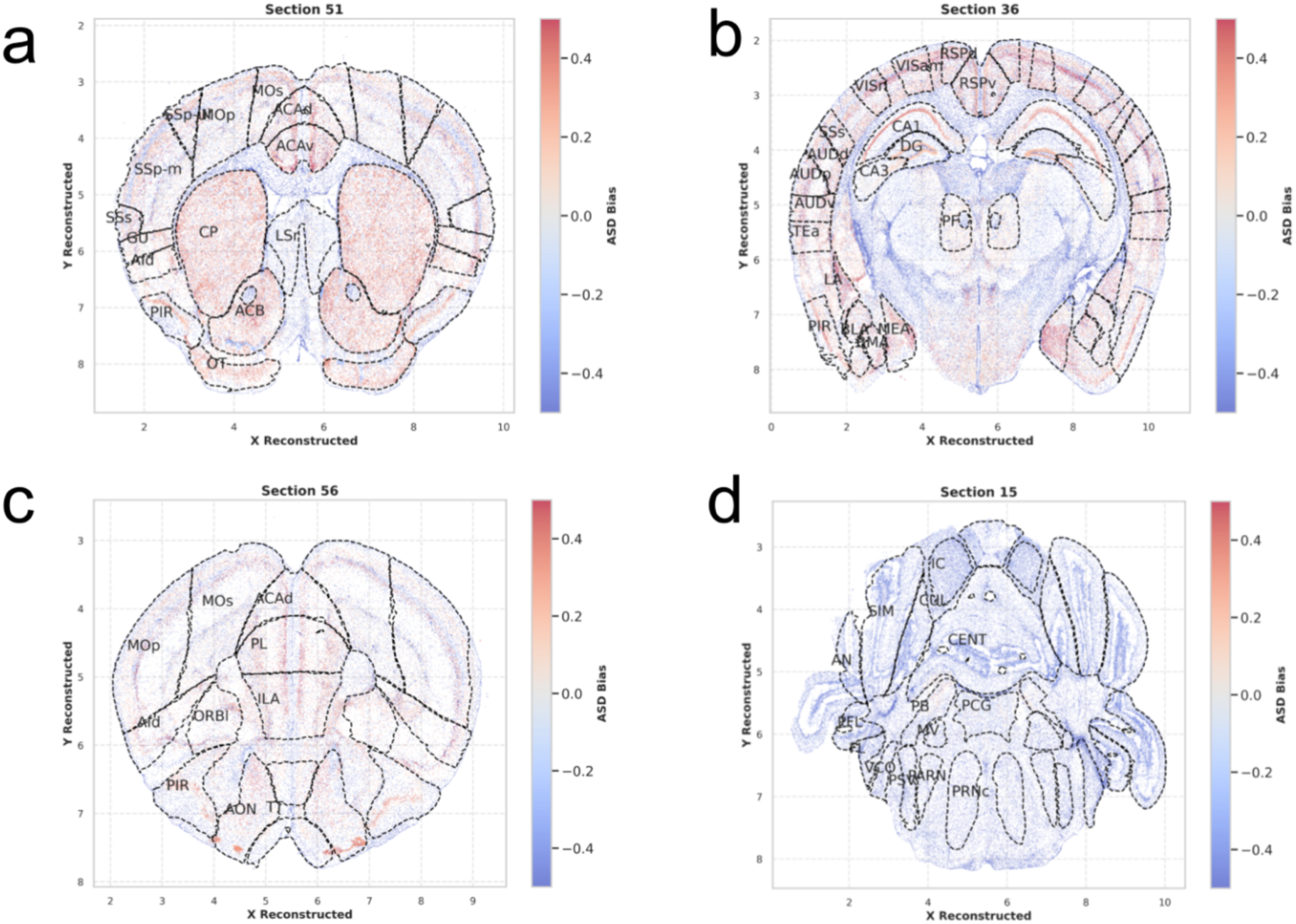
Single-cell spatial ASD mutation biases. Examples of coronal sections of the mouse brain. The X and Y axes show reconstructed Allen CCF coordinates and each point represents a single cell profiled by the MERFISH analysis. Point colors indicate the magnitude of ASD mutation bias towards the corresponding cell type cluster. **a.)** The coronal section 51 with the striatum: CP (Caudoputamen), ACB (Nucleus accumbens), LSr (Lateral septal nucleus rostral rostroventral part); olfactory area: PIR (Piriform area), OT (Olfactory tubercle); somatosensory: SSs (Supplemental somatosensory area), SSp-m (Primary somatosensory area mouth), SSp-ul (Primary somatosensory area upper limb); GU (Gustatory areas); and motor cortex: Mos (Secondary motor area), Mop (Primary motor area). **b.)** The coronal section 36 with the amygdala: BLA (Basolateral amygdalar nucleus), BMA (Basomedial amygdalar nucleus), LA (Lateral amygdalar nucleus); hippocampus DG (Dentate gyrus), CA1 (Field CA1), CA3 (Field CA3); the thalamus: PF (Parafascicular nucleus); visual cortex: VISrl (Posterior parietal association areas), VISam (Anteromedial visual area); auditory cortex: AUDv (Ventral auditory area), AUDp (Primary auditory area), AUDd (Dorsal auditory area). **c.)** The coronal section 56 with the prefrontal cortical areas: PL (Prelimbic area), ILA (Infralimbic area), ACAd (Anterior cingulate area dorsal part), ORBl (Orbital area lateral part). **d.)** The coronal section 15 with the cerebellum: CUL (Culmen), CENT (Central lobule), AN (Ansiform lobule), SIM (Simple lobule). The complete list of structure abbreviations is listed in Supplementary Table 1.

**Supplementary Figure 12:**
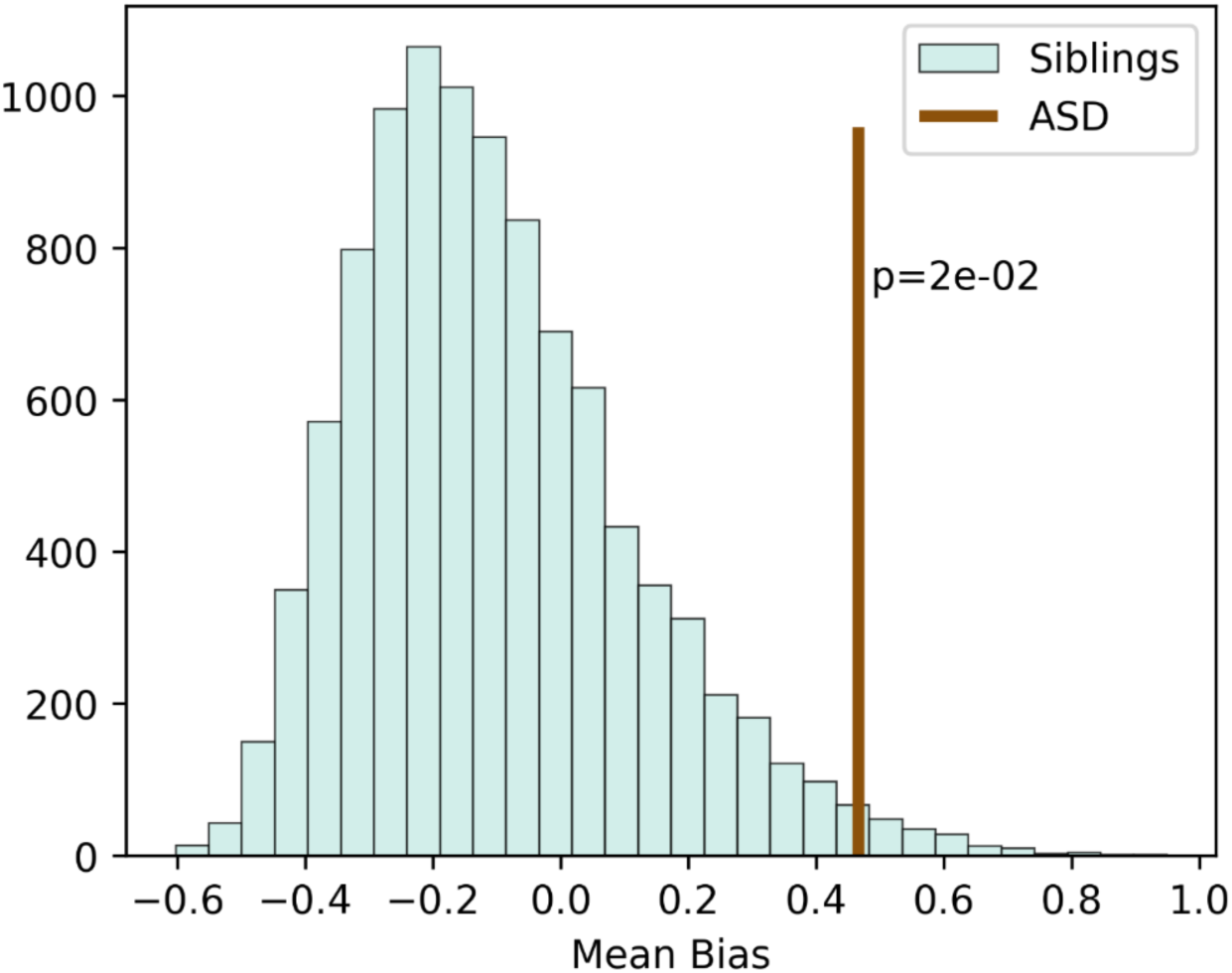
Enrichment of oxytocin receptor expression among the ASD circuit structures. The average gene expression bias (see Methods) for the oxytocin receptor gene, OXTR, across the 46 ASD circuit structures is represented by the brown vertical line. The distribution of the average OXTR gene expression biases for siblings is shown in cyan. To calculate the distribution for siblings, the genes with mutations in unaffected siblings were randomly subsampled to match the number of genes in the ASD gene set (60) and the 46 brain structures with the strongest mutation biases were selected for each of the 10,000 subsampled sibling sets. The distribution of the average OXTR gene expression biases for siblings was then calculated for these sets of structures.

## SUPPLEMENTARY TABLES

**Supplementary Table 1: ASD mutation biases across all mouse brain structures.** Comprehensive analysis of mutation biases calculated for 213 brain structures using the Allen Mouse Brain Atlas ISH data. For each structure, the table presents ASD mutation biases calculated using *de novo* mutations (LGD and Dmis) identified in SPARK exome-wide significant genes. Bias values reflect the weighted average of gene-specific expression biases, accounting for both mutation frequency and class enrichment. Statistical significance for each structure is determined by comparison with mutation biases calculated from 10,000 subsampled sibling gene sets.

**Supplementary Table 2: Connectome information score matrix for mouse brain structure connections.** Matrix of connectome weights representing the information content (–log probability) of connections between brain structure pairs, derived from the Allen Mouse Brain Connectivity Atlas. Connection probabilities are calculated by binning structure pairs based on their Cartesian distances and determining the likelihood of connections at each distance range. These scores are used to evaluate the significance of circuit connectivity in the GENCIC analysis by quantifying how surprising each connection is given the physical distance between structures.

**Supplementary Table 3: ASD mutation biases for cell types calculated using ABC atlas single-cell data.** Analysis of mutation biases at the cell type cluster level based on the Allen Brain Cell (ABC) atlas 10x v3 RNA sequencing data. For each cell type cluster, the table presents the ASD mutation bias calculated as a weighted average of gene-specific biases, accounting for both mutation frequency and class (LGD/Dmis). Gene expression specificity is determined by comparing against expression-matched random genes and weighted by v2-v3 correlation to account for technical variation. Statistical significance is assessed by comparison with mutation biases calculated from subsampled sibling gene sets.

**Supplementary Table 4: Cellular composition of brain structures within the ASD circuit.** Analysis of cell type distributions across circuit structures based on the Allen Brain Cell (ABC) atlas MERFISH data. For each circuit structure, the table presents the relative abundance of different cell types at the subclass level, with cell proportions indicated (proportions sum to 1 within each structure). Only cell types with robust representation (≥500 cells per structure) are included.

**Supplementary Table 5: Analysis of ASD mutation biases across circuit structures stratified by proband cognitive function.** Comparison of structure-specific mutation biases between two cohorts: probands with higher cognitive function (FSIQ > 70) and those with lower cognitive function (FSIQ ≤ 70). The table presents mutation biases calculated independently for each cohort using cohort-specific mutation weights, demonstrating how genetic impacts on different circuit structures relate to cognitive phenotypes. Statistical significance of bias differences between the two cohorts based on IQ-mutation permutation test is indicated for each structure.

## Notes

### Competing Interest Statement

The authors have declared no competing interest.

### Summary of Updates

No manuscript content change, only changed subject of area.

### Funding

- National Institute of Mental Health: NIMHR01MH124923

## References

1 Kanner, L. Autistic disturbances of affective contact. Acta Paedopsychiatr 35, 100–136 (1943).

2 Abrahams, B. S. & Geschwind, D. H. Advances in autism genetics: on the threshold of a new neurobiology. Nat Rev Genet 9, 341–355 (2008). 10.1038/nrg2346

3 Iossifov, I. et al. The contribution of de novo coding mutations to autism spectrum disorder. Nature 515, 216–221 (2014). 10.1038/nature13908

4 Satterstrom, F. K. et al. Large-Scale Exome Sequencing Study Implicates Both Developmental and Functional Changes in the Neurobiology of Autism. Cell 180, 568–584 e523 (2020). 10.1016/j.cell.2019.12.036

5 Zhou, X. et al. Integrating de novo and inherited variants in 42,607 autism cases identifies mutations in new moderate-risk genes. Nat Genet 54, 1305–1319 (2022). 10.1038/s41588-022-01148-2

6 Iakoucheva, L. M., Muotri, A. R. & Sebat, J. Getting to the Cores of Autism. Cell 178, 1287–1298 (2019). 10.1016/j.cell.2019.07.037

7 Jamain, S. et al. Mutations of the X-linked genes encoding neuroligins NLGN3 and NLGN4 are associated with autism. Nat Genet 34, 27–29 (2003). 10.1038/ng1136

8 Willsey, A. J. et al. Coexpression networks implicate human midfetal deep cortical projection neurons in the pathogenesis of autism. Cell 155, 997–1007 (2013). 10.1016/j.cell.2013.10.020

9 Gilman, S. R. et al. Rare de novo variants associated with autism implicate a large functional network of genes involved in formation and function of synapses. Neuron 70, 898–907 (2011). 10.1016/j.neuron.2011.05.021

10 Pinto, D. et al. Functional impact of global rare copy number variation in autism spectrum disorders. Nature 466, 368–372 (2010). 10.1038/nature09146

11 De Rubeis, S. et al. Synaptic, transcriptional and chromatin genes disrupted in autism. Nature 515, 209–215 (2014). 10.1038/nature13772

12 Parikshak, N. N. et al. Integrative functional genomic analyses implicate specific molecular pathways and circuits in autism. Cell 155, 1008–1021 (2013). 10.1016/j.cell.2013.10.031

13 Chang, J., Gilman, S. R., Chiang, A. H., Sanders, S. J. & Vitkup, D. Genotype to phenotype relationships in autism spectrum disorders. Nat Neurosci 18, 191–198 (2015). 10.1038/nn.3907

14 Sahin, M. & Sur, M. Genes, circuits, and precision therapies for autism and related neurodevelopmental disorders. Science 350 (2015). 10.1126/science.aab3897

15 Avena-Koenigsberger, A., Misic, B. & Sporns, O. Communication dynamics in complex brain networks. Nat Rev Neurosci 19, 17–33 (2017). 10.1038/nrn.2017.149

16 Alexander, G. E., DeLong, M. R. & Strick, P. L. Parallel organization of functionally segregated circuits linking basal ganglia and cortex. Annu Rev Neurosci 9, 357–381 (1986). 10.1146/annurev.ne.09.030186.002041

17 Rothwell, P. E. et al. Autism-associated neuroligin-3 mutations commonly impair striatal circuits to boost repetitive behaviors. Cell 158, 198–212 (2014). 10.1016/j.cell.2014.04.045

18 Zhou, Y. et al. Mice with Shank3 Mutations Associated with ASD and Schizophrenia Display Both Shared and Distinct Defects. Neuron 89, 147–162 (2016). 10.1016/j.neuron.2015.11.023

19 Menon, V. Large-scale brain networks and psychopathology: a unifying triple network model. Trends Cogn Sci 15, 483–506 (2011). 10.1016/j.tics.2011.08.003

20 Buch, A. M. et al. Molecular and network-level mechanisms explaining individual differences in autism spectrum disorder. Nat Neurosci 26, 650–663 (2023). 10.1038/s41593-023-01259-x

21 Lein, E. S. et al. Genome-wide atlas of gene expression in the adult mouse brain. Nature 445, 168–176 (2007). 10.1038/nature05453

22 Oh, S. W. et al. A mesoscale connectome of the mouse brain. Nature 508, 207–214 (2014). 10.1038/nature13186

23 Yao, Z. et al. A high-resolution transcriptomic and spatial atlas of cell types in the whole mouse brain. Nature 624, 317–332 (2023). 10.1038/s41586-023-06812-z

24 Golden, C. E., Buxbaum, J. D. & De Rubeis, S. Disrupted circuits in mouse models of autism spectrum disorder and intellectual disability. Curr Opin Neurobiol 48, 106–112 (2018). 10.1016/j.conb.2017.11.006

25 Gogos, J. A., Crabtree, G. & Diamantopoulou, A. The abiding relevance of mouse models of rare mutations to psychiatric neuroscience and therapeutics. Schizophr Res 217, 37–51 (2020). 10.1016/j.schres.2019.03.018

26 Zhang, M. et al. Molecularly defined and spatially resolved cell atlas of the whole mouse brain. Nature 624, 343–354 (2023). 10.1038/s41586-023-06808-9

27 Bult, C. J. et al. Mouse Genome Database (MGD) 2019. Nucleic Acids Res 47, D801–D806 (2019). 10.1093/nar/gky1056

28 Gandal, M. J. et al. Broad transcriptomic dysregulation occurs across the cerebral cortex in ASD. Nature 611, 532–539 (2022). 10.1038/s41586-022-05377-7

29 Jacquemont, S. et al. A higher mutational burden in females supports a “female protective model” in neurodevelopmental disorders. Am J Hum Genet 94, 415–425 (2014). 10.1016/j.ajhg.2014.02.001

30 Miettinen, K. Nonlinear multiobjective optimization. (Kluwer Academic Publishers, 1999).

31 Robertson, C. E. & Baron-Cohen, S. Sensory perception in autism. Nat Rev Neurosci 18, 671–684 (2017). 10.1038/nrn.2017.112

32 Euston, D. R., Gruber, A. J. & McNaughton, B. L. The role of medial prefrontal cortex in memory and decision making. Neuron 76, 1057–1070 (2012). 10.1016/j.neuron.2012.12.002

33 Rudebeck, P. H. & Murray, E. A. The orbitofrontal oracle: cortical mechanisms for the prediction and evaluation of specific behavioral outcomes. Neuron 84, 1143–1156 (2014). 10.1016/j.neuron.2014.10.049

34 Bush, G., Luu, P. & Posner, M. I. Cognitive and emotional influences in anterior cingulate cortex. Trends Cogn Sci 4, 215–222 (2000). 10.1016/s1364-6613(00)01483-2

35 Gogolla, N. The insular cortex. Curr Biol 27, R580–R586 (2017). 10.1016/j.cub.2017.05.010

36 Brown, S. P. et al. New Breakthroughs in Understanding the Role of Functional Interactions between the Neocortex and the Claustrum. J Neurosci 37, 10877–10881 (2017). 10.1523/JNEUROSCI.1837-17.2017

37 Leekam, S. R., Prior, M. R. & Uljarevic, M. Restricted and repetitive behaviors in autism spectrum disorders: a review of research in the last decade. Psychol Bull 137, 562–593 (2011). 10.1037/a0023341

38 Floresco, S. B. The nucleus accumbens: an interface between cognition, emotion, and action. Annu Rev Psychol 66, 25–52 (2015). 10.1146/annurev-psych-010213-115159

39 Wirtshafter, H. S. & Wilson, M. A. Lateral septum as a nexus for mood, motivation, and movement. Neurosci Biobehav Rev 126, 544–559 (2021). 10.1016/j.neubiorev.2021.03.029

40 Bhat, A. N., Landa, R. J. & Galloway, J. C. Current perspectives on motor functioning in infants, children, and adults with autism spectrum disorders. Phys Ther 91, 1116–1129 (2011). 10.2522/ptj.20100294

41 Peca, J. et al. Shank3 mutant mice display autistic-like behaviours and striatal dysfunction. Nature 472, 437–442 (2011). 10.1038/nature09965

42 Vertes, R. P., Linley, S. B. & Hoover, W. B. Limbic circuitry of the midline thalamus. Neurosci Biobehav Rev 54, 89–107 (2015). 10.1016/j.neubiorev.2015.01.014

43 Acsady, L. The thalamic paradox. Nat Neurosci 20, 901–902 (2017). 10.1038/nn.4583

44 Mitchell, A. S. & Chakraborty, S. What does the mediodorsal thalamus do? Front Syst Neurosci 7, 37 (2013). 10.3389/fnsys.2013.00037

45 Parnaudeau, S. et al. Mediodorsal thalamus hypofunction impairs flexible goal-directed behavior. Biol Psychiatry 77, 445–453 (2015). 10.1016/j.biopsych.2014.03.020

46 Allen, A. E., Procyk, C. A., Howarth, M., Walmsley, L. & Brown, T. M. Visual input to the mouse lateral posterior and posterior thalamic nuclei: photoreceptive origins and retinotopic order. J Physiol 594, 1911–1929 (2016). 10.1113/JP271707

47 Yang, C. et al. Medial prefrontal cortex and anteromedial thalamus interaction regulates goal-directed behavior and dopaminergic neuron activity. Nat Commun 13, 1386 (2022). 10.1038/s41467-022-28892-7

48 Cassel, J. C. et al. The reuniens and rhomboid nuclei of the thalamus: A crossroads for cognition-relevant information processing? Neurosci Biobehav Rev 126, 338–360 (2021). 10.1016/j.neubiorev.2021.03.023

49 Cai, R. Y., Richdale, A. L., Uljarevic, M., Dissanayake, C. & Samson, A. C. Emotion regulation in autism spectrum disorder: Where we are and where we need to go. Autism Res 11, 962–978 (2018). 10.1002/aur.1968

50 Banker, S. M., Gu, X., Schiller, D. & Foss-Feig, J. H. Hippocampal contributions to social and cognitive deficits in autism spectrum disorder. Trends Neurosci 44, 793–807 (2021). 10.1016/j.tins.2021.08.005

51 Phelps, E. A. Human emotion and memory: interactions of the amygdala and hippocampal complex. Curr Opin Neurobiol 14, 198–202 (2004). 10.1016/j.conb.2004.03.015

52 O’Mara, S. The subiculum: what it does, what it might do, and what neuroanatomy has yet to tell us. J Anat 207, 271–282 (2005). 10.1111/j.1469-7580.2005.00446.x

53 Janak, P. H. & Tye, K. M. From circuits to behaviour in the amygdala. Nature 517, 284–292 (2015). 10.1038/nature14188

54 Lebow, M. A. & Chen, A. Overshadowed by the amygdala: the bed nucleus of the stria terminalis emerges as key to psychiatric disorders. Mol Psychiatry 21, 450–463 (2016). 10.1038/mp.2016.1

55 Adhikari, A. et al. Basomedial amygdala mediates top-down control of anxiety and fear. Nature 527, 179–185 (2015). 10.1038/nature15698

56 Sias, A. C. et al. Dopamine projections to the basolateral amygdala drive the encoding of identity-specific reward memories. Nat Neurosci 27, 728–736 (2024). 10.1038/s41593-024-01586-7

57 Rolland, T. et al. Phenotypic effects of genetic variants associated with autism. Nat Med 29, 1671–1680 (2023). 10.1038/s41591-023-02408-2

58 Myers, S. M. et al. Insufficient Evidence for “Autism-Specific” Genes. Am J Hum Genet 106, 587–595 (2020). 10.1016/j.ajhg.2020.04.004

59 Buxbaum, J. D. et al. Not All Autism Genes Are Created Equal: A Response to Myers et al. Am J Hum Genet 107, 1000–1003 (2020). 10.1016/j.ajhg.2020.09.013

60 Sigurdsson, T. & Duvarci, S. Hippocampal-Prefrontal Interactions in Cognition, Behavior and Psychiatric Disease. Front Syst Neurosci 9, 190 (2015). 10.3389/fnsys.2015.00190

61 Chiang, A. H., Chang, J., Wang, J. & Vitkup, D. Exons as units of phenotypic impact for truncating mutations in autism. Mol Psychiatry 26, 1685–1695 (2021). 10.1038/s41380-020-00876-3

62 Feldman, I., Rzhetsky, A. & Vitkup, D. Network properties of genes harboring inherited disease mutations. Proc Natl Acad Sci U S A 105, 4323–4328 (2008). 10.1073/pnas.0701722105

63 Scangos, K. W., State, M. W., Miller, A. H., Baker, J. T. & Williams, L. M. New and emerging approaches to treat psychiatric disorders. Nat Med 29, 317–333 (2023). 10.1038/s41591-022-02197-0

64 Insel, T. R. & Cuthbert, B. N. Medicine. Brain disorders? Precisely. Science 348, 499–500 (2015). 10.1126/science.aab2358

65 Jin, X. et al. In vivo Perturb-Seq reveals neuronal and glial abnormalities associated with autism risk genes. Science 370 (2020). 10.1126/science.aaz6063

66 Siletti, K. et al. Transcriptomic diversity of cell types across the adult human brain. Science 382, eadd7046 (2023). 10.1126/science.add7046

67 van den Heuvel, M. P. & Sporns, O. A cross-disorder connectome landscape of brain dysconnectivity. Nat Rev Neurosci 20, 435–446 (2019). 10.1038/s41583-019-0177-6

68 Segal, A. et al. Regional, circuit and network heterogeneity of brain abnormalities in psychiatric disorders. Nat Neurosci 26, 1613–1629 (2023). 10.1038/s41593-023-01404-6

69 Meyer-Lindenberg, A. & Tost, H. Neural mechanisms of social risk for psychiatric disorders. Nat Neurosci 15, 663–668 (2012). 10.1038/nn.3083

70 Meyer-Lindenberg, A., Domes, G., Kirsch, P. & Heinrichs, M. Oxytocin and vasopressin in the human brain: social neuropeptides for translational medicine. Nat Rev Neurosci 12, 524–538 (2011). 10.1038/nrn3044

71 Leichsenring, F., Steinert, C., Rabung, S. & Ioannidis, J. P. A. The efficacy of psychotherapies and pharmacotherapies for mental disorders in adults: an umbrella review and meta-analytic evaluation of recent meta-analyses. World Psychiatry 21, 133–145 (2022). 10.1002/wps.20941

72 Porto, P. R. et al. Does cognitive behavioral therapy change the brain? A systematic review of neuroimaging in anxiety disorders. J Neuropsychiatry Clin Neurosci 21, 114–125 (2009). 10.1176/jnp.2009.21.2.114

